# The Gq/11 family of Gα subunits is necessary and sufficient for lower jaw development

**DOI:** 10.1101/2024.09.17.611698

**Authors:** Stanley M. Kanai, Chloe R. Garcia, MaCalia R. Augustus, Shujan A. Sharafeldeen, Elliott P. Brooks, Juliana Sucharov, Ezra S. Lencer, James T. Nichols, David E. Clouthier

**Affiliations:** Department of Craniofacial Biology, School of Dental Medicine, University of Colorado Anschutz Medical Campus, Aurora, CO USA; Department of Biology, Lafayette College, Easton, PA USA

**Keywords:** heterotrimeric G protein, craniofacial development, CRISPR/Cas9, single cell RNA-sequencing, small molecule inhibitor, zebrafish

## Abstract

Vertebrate jaw development is coordinated by highly conserved ligand-receptor systems such as the peptide ligand Endothelin 1 (Edn1) and Endothelin receptor type A (Ednra), which are required for patterning of lower jaw structures. The Edn1/Ednra signaling pathway establishes the identity of lower jaw progenitor cells by regulating expression of numerous patterning genes, but the intracellular signaling mechanisms linking receptor activation to gene regulation remain poorly understood. As a first step towards elucidating this mechanism, we examined the function of the Gq/11 family of Gα subunits in zebrafish using pharmacological inhibition and genetic ablation of Gq/11 activity and transgenic induction of a constitutively active Gq protein in *edn1^-/-^* embryos. Genetic loss of Gq/11 activity fully recapitulated the *edn1^-/-^* phenotype, with genes encoding G11 being most essential. Furthermore, inducing Gq activity in *edn1^-/-^*embryos not only restored Edn1/Ednra-dependent jaw structures and gene expression signatures but also caused homeosis of the upper jaw structure into a lower jaw-like structure. These results indicate that Gq/11 is necessary and sufficient to mediate the lower jaw patterning mechanism for Ednra in zebrafish.

**Summary statement:** Gq/11 is the signaling mediator downstream of Endothelin 1 and Endothelin Receptor Type A that drives tissue patterning for all lower jaw structures in zebrafish.

## Introduction

Endothelin A receptor (Ednra) and peptide ligand Endothelin-1 (Edn1) are essential for patterning the embryonic tissue that gives rise to lower jaw structures (Clouthier et al., 1998; Yanagisawa et al., 1998b; Miller et al., 2000; Ozeki et al., 2004). During development, Ednra is expressed in migratory and post-migratory cranial neural crest cells (NCCs) (Kurihara et al., 1995; Clouthier et al., 1998; Yanagisawa et al., 1998a; Miller et al., 2000; Nair et al., 2007; Ruest and Clouthier, 2009), which are multipotent progenitors that give rise to all bone and cartilage structures of the face (Le Douarin and Kalcheim, 1999; Chai et al., 2000; Tang and Bronner, 2020). Post-migratory cranial NCCs in pharyngeal arch one (or the mandibular portion of arch one in mammals) are stimulated by Edn1 that is secreted by the adjacent ventral ectoderm (Clouthier et al., 1998; Yanagisawa et al., 1998a; Miller et al., 2000), inducing gene expression that establishes two distinct patterning domains for lower jaw structures – a ventral patterning domain that gives rise to the mandible and Meckel’s cartilage and an intermediate patterning domain that gives rise to the jaw joint in zebrafish and middle ear structures in mammals (Clouthier et al., 2000; Miller et al., 2003; Ozeki et al., 2004; Jeong et al., 2008; Ruest and Clouthier, 2009; Talbot et al., 2010; Barske et al., 2016; Askary et al., 2017; Tavares et al., 2017). Edn1/Ednra also antagonizes the effects of Nr2f nuclear receptors and Jagged/Notch, two factors that establish upper jaw identity of cranial NCCs in the dorsal domain of the first and second arches (Zuniga et al., 2010; Barske et al., 2016; Teng et al., 2017; Barske et al., 2018). Thus, Edn1/Ednra plays the dual role of establishing lower jaw identity of cranial NCCs while also establishing the boundary separating the lower jaw and upper jaw patterning domains.

In mice and zebrafish, attenuation of the Edn1/Ednra signaling pathway diminishes intermediate and ventral patterning gene expression, causing expansion of dorsal patterning gene expression into the intermediate and ventral domains. This results in a homeotic transformation of lower jaw structures into upper jaw-like structures (Miller et al., 2000; Ozeki et al., 2004; Ruest and Clouthier, 2009; Pritchard et al., 2020; Kanai et al., 2022). Conversely, ectopic activation of the Edn1/Ednra signaling pathway in the dorsal patterning domain (or the maxillary portion of arch one in mammals) results in homeotic transformation of upper jaw structures into lower jaw-like structures (Sato et al., 2008b; Alexander et al., 2011; Zuniga et al., 2011; Tavares and Clouthier, 2015; Barske et al., 2018; Kurihara et al., 2023). In either case where Edn1/Ednra signaling is lost or ectopically activated, misexpression of patterning genes that are positively or negatively regulated by Edn1/Ednra changes the positional identities of cranial NCCs. While the Edn1/Ednra-regulated genes have been extensively characterized, the intracellular signaling events connecting Ednra activation to gene regulation are not fully understood.

Ednra signaling is mediated by heterotrimeric G proteins, a complex composed of a Gα subunit and a Gβ/Gγ obligate heterodimer (Gβγ) (Gilman, 1987). Ligand-bound Ednra facilitates the exchange of GDP for GTP on the Gα subunit, resulting in Gα and Gβγ to dissociate and interact with their respective signaling effectors (Oldham and Hamm, 2008). Ednra can couple with all four Gα subunit family members, Gq/11, G12/13, Gs and Gi/o, to varying extents depending on the cell and tissue type (Aramori and Nakanishi, 1992; Khac et al., 1994; Kawanabe et al., 2002; Wettschureck and Offermanns, 2005; Inoue et al., 2019; Okashah et al., 2019; Masuho et al., 2023). In mice, Gq and G11 proteins are encoded by *Gnaq* and *Gna11,* respectively. Removal of *Gnaq* and *Gna11* using conventional knockout alleles results in midgestational lethality prior to the development of craniofacial structures, though embryos with one wild-type allele for *Gnaq* (*Gnaq^+/-^; Gna11^-/-^)* or *Gna11 (Gnaq^-/-^;Gna11^+/-^)* can survive to term (Offermanns et al., 1998). Notably, *Gnaq^-/-^;Gna11^+/-^*embryos exhibit defects that appeared to be confined to the jaw joint and middle ear structures, while *Gnaq^+/-^;Gna11^-/-^* embryos displayed no craniofacial defects (Offermanns et al., 1998). This suggests that while both *Gnaq* and *Gna11* are required for early development, *Gnaq* is specifically required for patterning the intermediate domain. Similarly, embryos with a combination of conventional knockout alleles for *Gna11* and NCC-specific conditional knockout alleles for *Gnaq* (*Gnaq^flox/flox^;Gna11^-/-^;*P0-Cre) exhibited craniofacial defects that appeared to be confined to proximal lower jaw structures, including fusion of the jaw joint and hypoplasia of the proximal mandible, tympanic ring, malleus and incus (Dettlaff-Swiercz et al., 2005). In contrast, the entire mandible undergoes a homeotic transformation into a maxilla-like structure in mice lacking *Edn1*, *Ednra or Ece1,* a gene that encodes an essential enzyme in the Edn1 biosynthetic pathway (Yanagisawa et al., 1998b; Ozeki et al., 2004; Ruest and Clouthier, 2009). These differences suggest that there are Gq/11 dependent and independent sub-domains within the mandibular arch, where patterning of the Gq/11 independent domain is mediated by a different Gα family member (Sato et al., 2008a). Arguing against this model, mice homozygous for knockout alleles of Gs, G12/13 or Gi/o do not exhibit patterning defects in distal lower jaw structures (Jiang et al., 1998; Dettlaff-Swiercz et al., 2005; Wettschureck and Offermanns, 2005; Plummer et al., 2012; Lei et al., 2016). Thus, the question remains whether patterning is mediated solely by Gq/11 or a combination of Gq/11 and other Gα family members

This question has direct significance for human health. Perturbations to the Ednra signaling pathway are the underlying causes of congenital syndromic disorders such as Mandibulofacial dysostosis with alopecia (Gordon et al., 2015; Kurihara et al., 2023), Oro-oto-cardiac syndrome (Pritchard et al., 2020) and Auriculocondylar Syndrome (Rieder et al., 2012; Gordon et al., 2013; Marivin et al., 2016; Kanai et al., 2022). Thus, clarification of the G protein family or families mediating Edn1/Ednra-dependent jaw patterning will help elucidate additional signaling pathway components that are linked to human craniofacial differences.

In this study, we examined the role of Gq/11 in craniofacial patterning using multiple orthogonal approaches in zebrafish. Using a Gq/11 small molecule inhibitor YM-254890 (YM) (Takasaki et al., 2004; Nishimura et al., 2010), we demonstrate that YM-treated zebrafish embryos exhibit craniofacial phenotypes and gene expression changes indicative of Ednra signaling attenuation. In addition, we created mutant alleles for three genes that encode zebrafish Gq/11 proteins, *gnaq*, *gna11a* and *gna11b*, and show that triple homozygous mutant larvae exhibit craniofacial phenotypes nearly identical to those observed in *edn1^-/-^* mutant larvae. Furthermore, the craniofacial phenotypes in *edn1^-/-^* mutant embryos can be rescued by inducing expression of a constitutively active form of Gq during craniofacial patterning. Taken together, these results suggest that Gq/11 is necessary and sufficient for establishing intermediate and ventral patterning domains and the subsequent development of all Ednra-dependent skeletal elements of the lower jaw in zebrafish.

## Results

### YM-254890 causes craniofacial phenotypes resembling partial loss of Edn1/Ednra signaling

To begin our analysis of the G protein signaling pathway downstream of Ednra in zebrafish craniofacial development, we examined the effect of treating zebrafish embryos with YM-254890 (YM), a naturally derived compound from *Chromobacterium* species that inhibits a subset of Gq/11 proteins with high selectivity and potency (Takasaki et al., 2004; Nishimura et al., 2010). YM binds to a hydrophobic cleft in mammalian Gq in a region spanning the GTPase (Ras-like) and α-helical domains, stabilizing the allosteric mechanism that promotes GDP release, the rate-limiting step for Gα activation (Nishimura et al., 2010; Flock et al., 2015). This inhibitory effect is dependent on a network of hydrophobic interactions between YM and eight key residues that are present in Gq, G11 and G14, but not in G15 or other Gα family members (Fig. 1) (Nishimura et al., 2010; Onken et al., 2018). In zebrafish, these eight residues are highly conserved in Gq, G11 and G14, weakly conserved in G15 and highly divergent in other Gα family members (Fig. 1), suggesting that YM should display highly specific inhibitory effects towards the Gq/11 signaling pathway in zebrafish, with the exception of G15. Because the expression of G15 is restricted to sensory cells (Ohmoto et al., 2011; Oka and Korsching, 2011), it is unlikely that G15 signaling activity would interfere with our analysis of craniofacial development.

**Figure 1.**
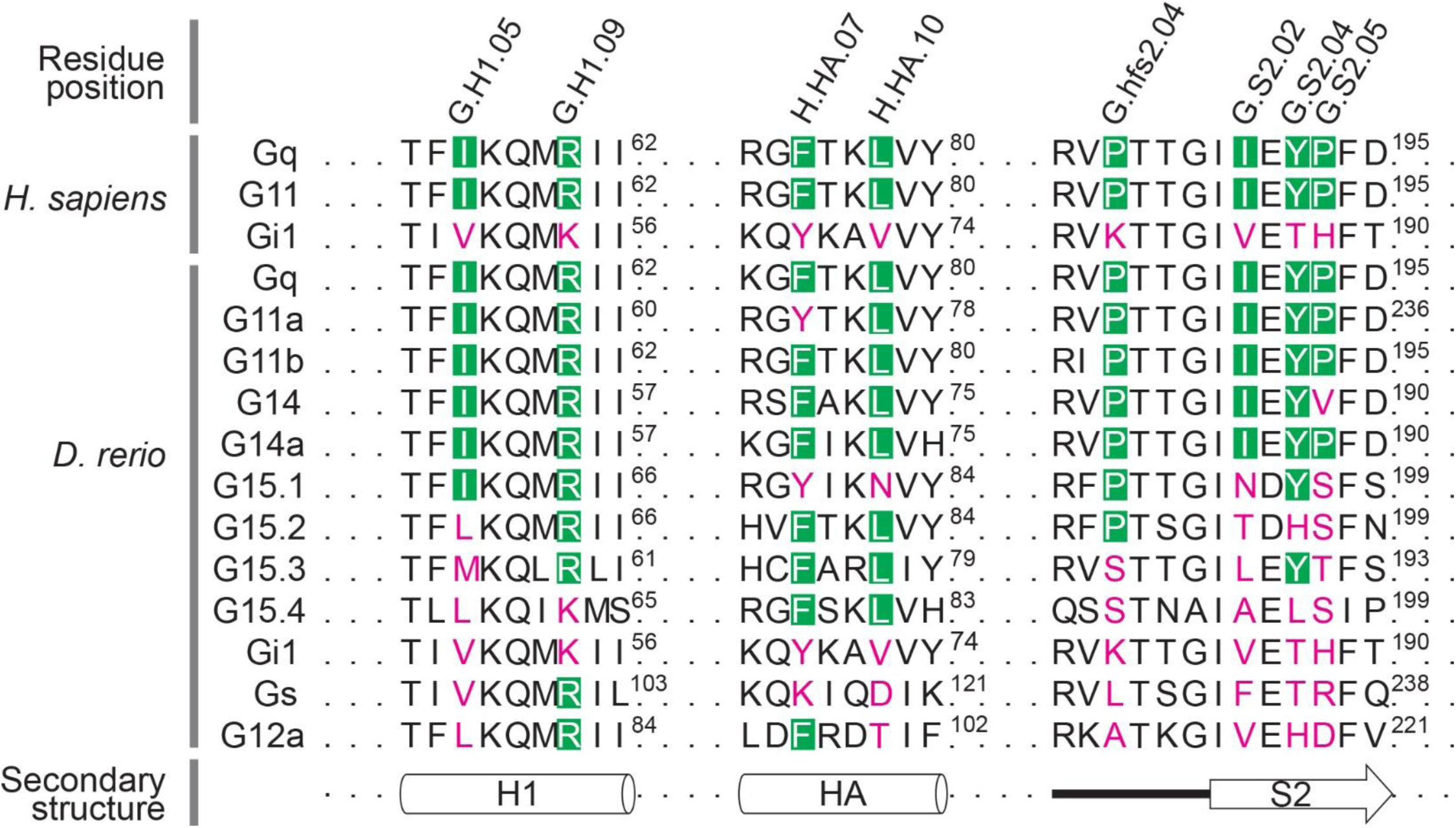
Sensitivity to YM is determined by eight conserved residues in a subset of Gq/11 family members. In zebrafish, Gq/11 family members are encoded by *gnaq* (Gq), *gna11a* (G11a), *gna11b* (G11b), *gna14* (G14), *gna14a* (G14a) and *gna15.1-15.4* (G15.1-15.4). Residue positions in Gq/11 family members that interact with YM are annotated with the Common Gα Numbering (CGN) system (Flock et al. 2015). YM-binding residues that are conserved in a subset of Gq/11 family members are shown in green boxes, while diverged residues are labeled in magenta. Superscript numbers indicate the amino acid position for the respective Gα protein. H1 and HA are α-helices of the Ras-like domain and the α helical domain, respectively, and S2 is a β-sheet of the Ras-like domain. These structures constitute the hydrophobic binding pocket for YM in a subset of Gq/11 family members.

To test whether YM treatment would produce craniofacial phenotypes, wild-type embryos were incubated with YM or solvent control [dimethyl sulfoxide (DMSO)] between 16-36 hours post fertilization (hpf), the approximate window for Edn1/Ednra-dependent craniofacial patterning (Miller et al., 2000; Miller et al., 2003; Walker et al., 2006; Kimmel et al., 2007; Vieux-Rochas et al., 2010; Alexander et al., 2011; Zuniga et al., 2011; Barske et al., 2016; Barske et al., 2018; Meinecke et al., 2018), and then grown to 6 days post fertilization (dpf) and processed for bone and cartilage staining. We tested a wide range of YM concentrations (from 1 nM to 100 uM), though we did not exceed 100 uM due to the embryonic toxicity threshold of the solvent, DMSO (Hoyberghs et al., 2021). Only 100 uM YM produced craniofacial phenotypes, which included an open mouth and uninflated swim bladder (Fig. 2B,D), all indications of defective jaw function (Neuhauss et al., 1996). Additional Edn1/Ednra pathway-associated defects were further visible in flat-mounted preparations of dissected viscerocrania, including hypoplasia of the palatoquadrate and symplectic cartilages, fusion of the jaw and hyomandibular joints and fusion of the opercle and branchiostegal ray dermal bones (Fig. 2F,I). Defects to at least one of these structures were found in 93% (26/28) of wild-type larvae treated with YM, with considerable variation in the laterality and number of skeletal elements affected per larvae (Fig. 2K).

**Figure 2.**
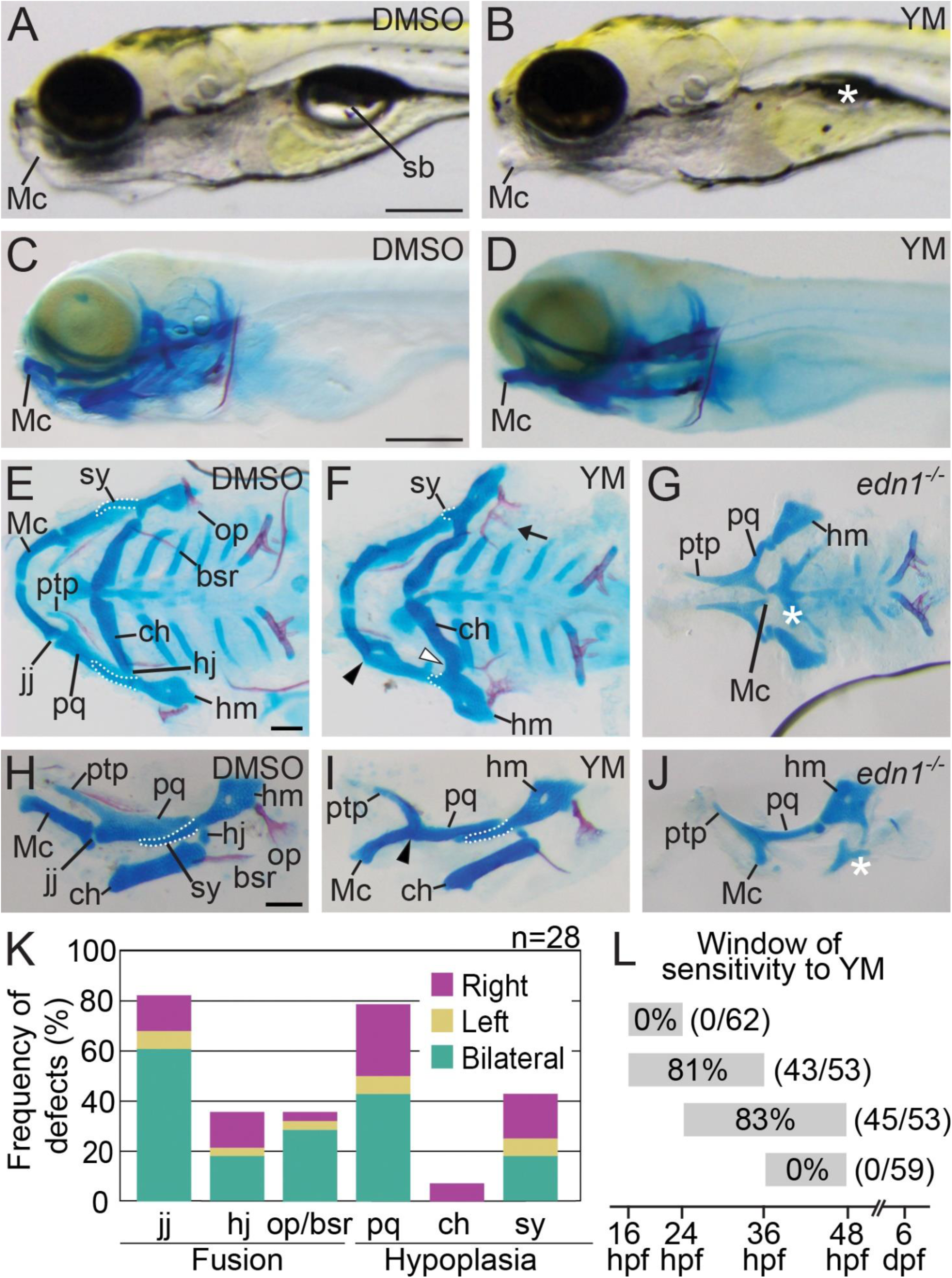
YM causes defects to a subset of Edn1/Ednra-dependent lower jaw skeletal elements. **(A-D)** Lateral views of 6 dpf embryos treated with DMSO (A) or YM (B), shown in gross (A,B) or whole-mount skeletal preparations of larvae (C,D). The asterisk in (B) indicates the absence of inflated swim bladder. **(E-J)** Ventral (E-G) or lateral (H-J) views of flat-mounts for the viscerocranium from 6 dpf wild-type larvae treated with DMSO (E,H) or YM (F,I), or an untreated *edn1^-/-^*larvae (G,J). In E,F,H,I, white outlines highlight the symplectic cartilage (sy). In F,I, black arrowheads indicate fusion of the jaw joint (jj), white arrowhead indicates fusion of the hyomandibular joint (hj) and the arrow indicates fusion of the opercle (op) and branchiostegal ray (bsr). In G,J, the white asterisk indicates absence of the ceratohyal (ch). **(K)** Frequency of defects in six Edn1/Ednra-dependent skeletal elements in larvae treated with YM from 16-36 hpf. Absence of structure, hypoplasia, or fusion were scored as defects (accounting for sidedness). A total of 28 larvae were examined across two independent experiments. The effect of YM was determined to be statistically significant with a chi-square test (*p*=0.0001), comparing the number of defects in DMSO-treated larvae (0) to YM-treated larvae. **(L)** A schematic illustrating the four overlapping time intervals of YM application (grey bars). Numbers within bars represent the percentages of larvae exhibiting at least one defect to a Edn1/Ednra-dependent lower jaw structure. The total number of larvae examined is given in parentheses. Scale bars are 500 μm (A,C) and 100 μm (E, H). hm; hyomandibular, Mc; Meckel’s cartilage, pq; palatoquadrate ptp; pterygoid process of the palatoquadrate.

Phenotypes resulting from YM treatment were not as severe as phenotypes caused by a complete loss of Edn1/Ednra signaling. Larvae homozygous for a new *edn1* allele lacking the majority of the coding sequence for the Edn1 bioactive peptide [distinct from the *sucker* allele that causes an amino acid substitution (Miller et al., 2000); Fig. S1] exhibited absence and severe hypoplasia of ventral structures (Fig. 2G,J). In contrast, the phenotypes of YM-treated larvae more closely resembled phenotypes caused by partial loss of Edn1/Ednra signaling (Walker et al., 2006; Kimmel et al., 2007; Miller et al., 2007; Talbot et al., 2010). These data suggest that YM-induced defects are caused by partial inhibition of the Edn1/Ednra signaling pathway.

To determine the developmental time during which Gq/11 activity is required for lower jaw development, embryos were treated with 100 uM YM across four overlapping time intervals (16-24, 16-36, 24-48 or 36-48 hpf) and then analyzed for craniofacial skeleton defects at 6 dpf. Craniofacial defects were observed only in embryos treated between 16-36 or 24-48 hpf (Fig. 2L), with the percentage of affected larvae (Fig. 2L) and frequency and severity of affected skeletal elements being similar for both time intervals (Fig. S2). These results suggest that Gq/11 activity is required for patterning between 24-36 hpf. This window of sensitivity to YM is similar to the window of sufficiency for Edn1/Ednra signaling in craniofacial patterning in zebrafish and mice (Miller et al., 2000; Miller et al., 2003; Walker et al., 2006; Kimmel et al., 2007; Ruest and Clouthier, 2009; Vieux-Rochas et al., 2010; Alexander et al., 2011; Zuniga et al., 2011; Barske et al., 2016; Barske et al., 2018; Meinecke et al., 2018).

### YM causes gene expression changes in cranial NCCs that resemble partial loss of Edn1/Ednra signaling

The loss of Edn1/Ednra signaling causes characteristic changes to gene expression in NCCs of the first and second pharyngeal arches, which include downregulation of lower jaw patterning genes and upregulation of upper jaw patterning genes in the intermediate and ventral domains (Clouthier et al., 1998; Thomas et al., 1998; Clouthier et al., 2000; Miller et al., 2000; Miller et al., 2003; Ozeki et al., 2004; Ruest et al., 2004; Walker et al., 2006; Nair et al., 2007; Walker et al., 2007; Sato et al., 2008a; Tavares et al., 2012; Barske et al., 2016; Askary et al., 2017; Tavares et al., 2017; Barske et al., 2018). To determine whether YM caused similar changes to gene expression in cranial NCCs as does partial loss of Edn1/Ednra signaling, we performed single cell RNA-sequencing (scRNA-seq) on *sox10:mRFP;fli1a:EGFP* double-positive cells from embryos treated with DMSO or 100 uM YM between 16-36 hours (Fig. 3A). Subsequent analyses with Seurat identified 12 cell clusters with unique transcriptional profiles (Fig. 3B, Fig. S3, Table S1). Using known marker genes (Miller et al., 2003; Askary et al., 2017; Barske et al., 2018; Mitchell et al., 2021; Fabian et al., 2022; Stenzel et al., 2022), NCC populations were identified from the frontonasal region (*akap12b+*, *alx4a+*), anterior pharyngeal arches 1 and 2 (*dlx2a+, dlx5a+*) and posterior pharyngeal arches 3-7 (*prdm1+*) (Fig. 3B, D). The NCCs of pharyngeal arches 1 and 2 were distributed across three clusters, with one cluster representing the ventral domain cells of both arches 1 (*hand2+*) and 2 (*hand2+*, *hoxb2a+*), and two separate clusters representing dorsal domain cells of arch 1 *(pou3f3b+*, Hox-) and arch 2 (*jag1b+*, *hoxb2a+*). Although we did not find a dedicated cluster for intermediate domain cells, we used known marker genes to identify the approximate population of intermediate domain cells nested within the ventral and dorsal clusters (Fig. 3C, Fig. S4). The remaining clusters appeared to consist of NCC-derived pigment cells (cluster 11: *dct*+, *tyrp1b*+) and hematopoietic derivatives (cluster 9: *hbbe3+, hbae3+*; cluster 10: *myb+,lyz+*); these clusters were not examined further in this study.

**Figure 3.**
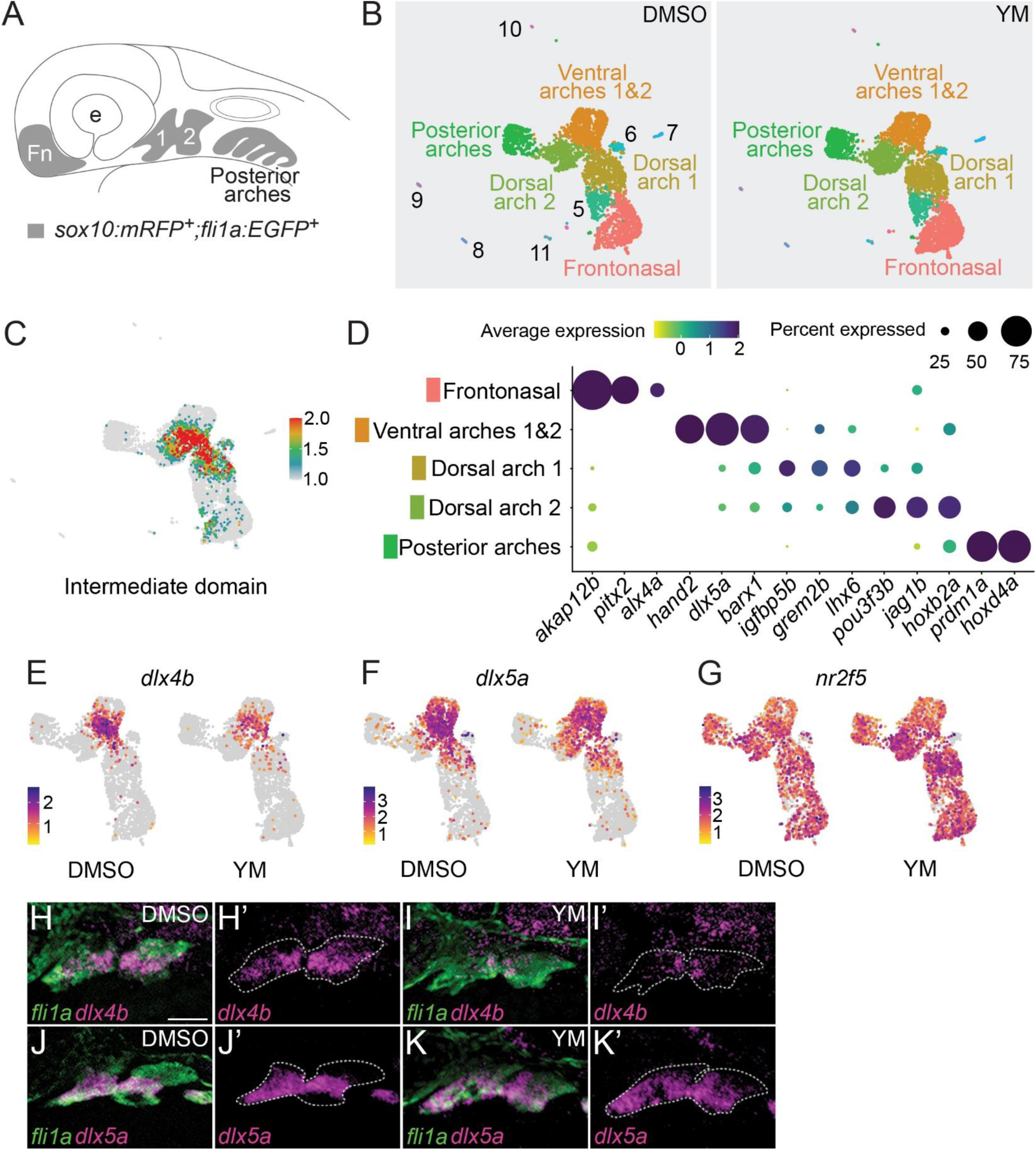
YM reduces expression of intermediate patterning genes and increases expression of dorsal patterning genes. **(A)** Schematic of a zebrafish embryo at 36 hpf. Cells double-labeled with *sox10:mRFP* and *fli1a:EGFP* transgenic reporters (highlighted in grey) represent cranial neural crest populations from the frontonasal region (Fn), anterior pharyngeal arches 1 (1) and 2 (2) and posterior pharyngeal arches. **(B)** UMAP plots for DMSO or YM-treated samples. Clusters analyzed in this study are labeled with the NCC populations they represent. Clusters 5-11 are described further in Supplementary Information (Fig. S3, Table S1). Equivalent clusters between DMSO and YM-treated samples are labeled with the same colors. **(C)** Feature map highlighting approximate cell populations in the intermediate domains of pharyngeal arches 1 and 2, shown overlayed on combined UMAP plots of control and YM-treated samples. The feature map represents the composite average expression level for 14 experimentally verified intermediate domain patterning genes, *ccn2b, dlx3b*, *dlx4a*, *dlx4b*, *emx2*, *fgfbp2a*, *foxc1b, foxd1, fsta, grem2b, igfbp5b, msx1a, nkx3.2* and *shox* (Fig. S4). The scale is average expression. **(D)** Dot plot of selected marker genes and their respective cluster identity (Fig. S3). **(E-G)** Feature maps highlighting differential expression of *dlx4b* (E), *dlx5a* (F) and *nr2f5* (G) in DMSO or YM-treated samples. The scale is average expression. **(H-K)** Fluorescence in situ hybridization and immunofluorescence of 36 hpf embryos treated with DMSO or YM between 16-36 hpf. DMSO (H,J) or YM-treated (I,K) embryos were probed for *dlx4b* (H,I) or *dlx5a* (J,K) using fluorescence in situ hybridization (magenta). Pharyngeal arches, labeled with *fli1a:EGFP*, were detected with immunofluorescence (green). Approximate borders for pharyngeal arches 1 and 2 are indicated with dashed lines (H’,I’,J’,K’). Images are representative of four embryos. Scale bar is 50 μm.

To examine the effects of YM on gene expression, we performed differential expression (DE) analysis between equivalent clusters in YM-treated samples relative to control samples (Table S2). DE genes with adjusted p values less than 0.05 were found in six out of eleven clusters – frontonasal, ventral arches 1 and 2, dorsal arch 1, dorsal arch 2, posterior arches 3-7 and cluster 5 (unknown identity) (Fig. S5, Table S2). Overall, a total of 36 and 34 unique, non-overlapping genes were upregulated or downregulated, respectively, in YM-treated samples. Of the aberrantly upregulated genes, dorsal patterning genes including *hey1, her6, her9* and *nr2f5* were present in the ventral and dorsal arch clusters (Fig. 3G, Fig. S5, Table S2) (Zuniga et al., 2010; Barske et al., 2016; Askary et al., 2017; Barske et al., 2018). The majority of aberrantly downregulated genes were in the ventral arches 1 and 2 and dorsal arch 1. In the ventral arches cluster, more than half of the reduced genes were Ednra targets. Notably, the intermediate patterning genes *dlx4a* and *dlx4b* (Fig. 3E,H,I, Fig. S5) were downregulated to a greater extent than ventral patterning genes like *hand2* and *dlx5a* (Fig. 3F,J,K, Fig. S5). These gene expression changes are similar to changes observed in animal models with partial loss of Edn1/Ednra signaling (Walker et al., 2006; Miller et al., 2007; Walker et al., 2007; Sato et al., 2008a; Ruest and Clouthier, 2009; Tavares et al., 2012). Furthermore, the relatively modest downregulation of ventral patterning genes compared with intermediate patterning is consistent with the unperturbed ventral domain-derived skeletal structures in YM-treated larvae (Fig. 2F,I). These results indicate that YM is partially attenuating the Edn1/Ednra signaling pathway in cranial NCCs through the inhibition of Gq/11.

### Pharmacogenetic interactions suggest that Gq/11 is also required for ventral domain patterning

Because we could not use a higher concentration of YM due to solvent (DMSO) toxicity, we could not distinguish whether the YM-generated phenotype was due to incomplete inhibition of Gq/11 activity or whether Gq/11 activity was only required in the intermediate domain. To address this question, we tested the effects of YM on *edn1^+/-^*zebrafish embryos, with the supposition that these embryos would have reduced expression of Ednra-dependent genes similar to *Edn1^+/-^* mouse embryos (Vieux-Rochas et al., 2010) and thus would be sensitized to the effects of YM. Embryos generated from *edn1^+/-^*mating pairs were treated with 100 uM YM or DMSO between 16-36 hpf, then processed for skeletal preparations at 6 dpf. Craniofacial defects were more severe in YM-treated *edn1^+/-^* larvae compared with YM-treated wild-type embryos (Fig. 4). We observed defects in ventral domain derivatives that were present in YM-treated *edn1^+/-^* larvae but not in YM-treated wild-type larvae, such as hypoplasia of the ventral portion of the Meckel’s cartilage and absence of the ceratohyal (Fig. 4B,E). The number of skeletal elements affected in individual larvae was variable in both YM and DMSO conditions, but overall, *edn1^+/-^* larvae had defects to more skeletal elements per individual compared to wild-type larvae (Fig. 4G). YM treatment had no effect on the phenotype of *edn1^-/-^* larvae (Fig. 4C,F), indicating that YM is specifically acting on Gq/11 downstream of Ednra and not on other Gq/11-couple receptors. The increased frequency of defects elicited by this pharmacogenetic interaction (Fig. 4G,H) suggests that YM only partially inhibits Gq/11 activity in wild-type zebrafish embryos, while a sensitized background like *edn1^+/-^* can augment the inhibitory effect of YM. Furthermore, the increased phenotype severity elicited by this pharmacogenetic interaction (Fig. 4D,E) suggests Gq/11 is required for patterning beyond the intermediate domain.

**Figure 4.**
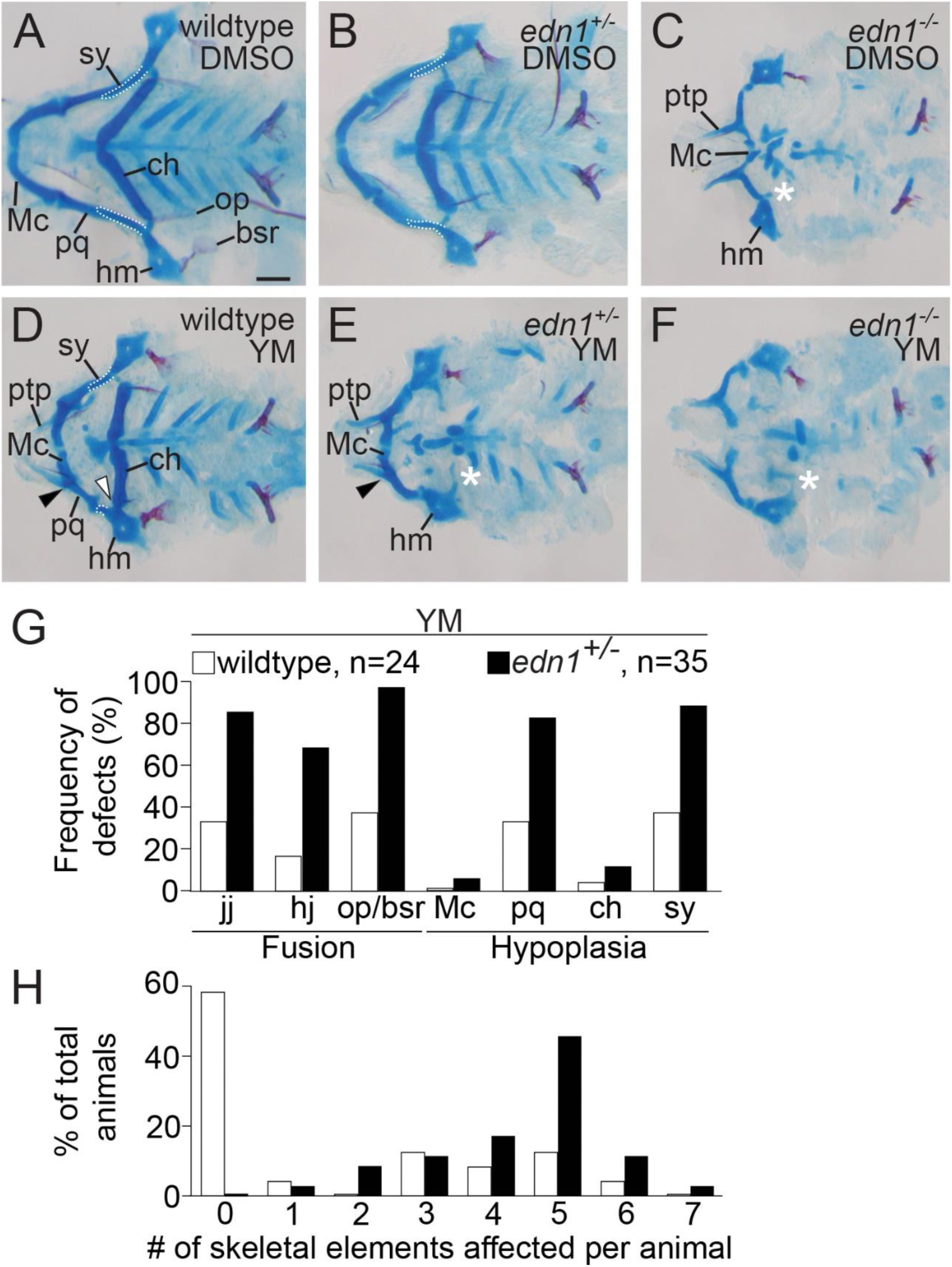
YM increases the prevalence and severity of lower jaw defects in *edn1^+/-^* larvae relative to wild-type alarvae. **(A-F)** Representative flat-mounts of viscerocranium from wild-type (A,D), *edn1^+/-^* (B,E), or *edn1^-/-^*(C,F) larvae treated with DMSO (A-C) or 100 μM YM (D-F) between 16-36 hpf. In A,B,D, white outlines highlight the symplectic cartilage. In C,E,F, the white asterisk indicate absence of the ceratohyal. In D,E, black arrowheads indicate fusion of the jaw joint. In D, the white arrowhead indicates fusion of the hyomandibular joint. All skeletal preparations are 6 dpf. **(G)** Frequency of defects in seven Edn1/Ednra-dependent skeletal elements in YM-treated wild-type or *edn1^+/-^*larvae. Absence of structure, hypoplasia, or fusion were scored as defects (ignoring sidedness). The pharmacogenetic interaction between *edn1* and YM was determined to be statistically significant with a chi-square test (*p*=0.0001), comparing the number of defects in YM-treated *edn1^+/+^* and *edn1^+/-^* larvae. **(H)** Percentage of individual larvae presenting with defects in the seven Edn1/Ednra-dependent structures. Individual larvae were scored for total number of skeletal elements affected (ignoring sidedness). Bar graphs are color-coded the same as G. All wild-type, *edn1^+/-^*or *edn1^-/-^* embryos treated with DMSO or YM are siblings from the same clutch. Scale bar is 100 μm.

### G11 paralogs are necessary for development of all Edn1/Ednra-dependent craniofacial structures

To more comprehensively interrogate the role of Gq/11 in facial development, we mutated genes encoding for Gq and G11 proteins using CRISPR/Cas9. In zebrafish, Gq is encoded by *gnaq,* and G11 is encoded by two paralogs *gna11a* and *gna11b.* Although *gnaq* expression was undetectable in our scRNA-seq analysis (Fig. S6), we proceeded to target the *gnaq* allele given the importance of *Gnaq* in mouse lower jaw development (Offermanns et al., 1998). Other Gq/11 family members, including *gna14, gna14a* and *gna15.1-15.4*, were not considered because they exhibit cell-type specific expression in sensory organs and hematopoietic lineages (Ohmoto et al., 2011; Oka and Korsching, 2011). We designed CRISPR/Cas9 guide RNAs to target exon 3 or 4 of the genes (Fig. S7A), similar to targeting strategies used for *Gnaq* and *Gna11* knockout alleles in mice (Offermanns et al., 1998). For all three genes, we identified a frame-shift mutation that is predicted to result in a premature stop codon and non-functional protein product (Fig. S7B). Similar to wild-type larvae (Fig. 1,2), larvae and adults that were homozygous for a single mutant allele (*gnaq^-/-^*, *gna11a^-/-^* or *gna11b^-/-^*) (data not shown) or triple heterozygous for all mutant alleles (*gnaq^+/-^;gna11a^+/-^;gna11b^+/-^*) (Fig. 5H) did not exhibit overt craniofacial phenotypes and were viable and fertile. However, we observed a spectrum of craniofacial phenotypes in larvae generated from *gnaq^+/-^;gna11a^+/-^;gna11b^+/-^* mating pairs (Fig. 5). A total of 336 larvae were processed for skeletal preparations at 6 dpf, followed by analysis of genotypes and phenotypes (Fig. S7). The most severe defects were observed in larvae that were *gnaq^+/+^;gna11a^-/-^;gna11b^-/-^*(Fig. S7C), *gnaq^+/-^ ;gna11a^-/-^;gna11b^-/-^*(Fig. 5B) or *gnaq^-/-^;gna11a^-/-^;gna11b^-/-^* (Fig. 5A), exhibiting defects in all Edn1/Ednra-dependent skeletal elements and resembling the *edn1^-/-^* phenotype (Fig. 2G,J). We then examined whether phenotype severity and penetrance were sensitive to gene dosage by quantifying the genotype-phenotype relationship for various allele combinations (Fig. 5I). Whereas all larvae with only one wild-type allele of *gnaq* (*gnaq^+/-^;gna11a^-/-^;gna11b^-/-^*) produced the *edn1^-/-^* phenotype, larvae with only one wild-type allele of *gna11a* (*gnaq^-/-^;gna11a^+/-^;gna11b^-/-^*) or *gna11b* (*gnaq^-/-^;gna11a^-/-^;gna11b^+/-^*) produced highly variable phenotypes that were all less severe than *edn1^-/-^*larvae (Fig. 5C,D). The addition of a wild-type *gnaq* allele to one wild-type allele of *gna11a* or *gna11b* (*gnaq^+/-^;gna11a^+/-^;gna11b^-/-^* and *gnaq^+/-^;gna11a^-/-^;gna11b^+/-^*) produced the same phenotype severity and variation as larvae with only one wild-type allele for *gna11a* or *gna11b* (Fig. 5E,F). However, larvae with a wild-type allele of *gna11a* and *gna11b* (*gnaq^-/-^;gna11a^+/-^;gna11b^+/-^*) exhibited reduced phenotype severity and variation compared with larvae having only one wild-type allele of *gna11a* or *gna11b* (Fig. 5G). Taken together, these results suggest that *gna11a* and *gna11b* are necessary for the development of Edn1/Ednra-dependent craniofacial structures in a gene-dosage dependent manner, while *gnaq* appears dispensable.

**Figure 5.**
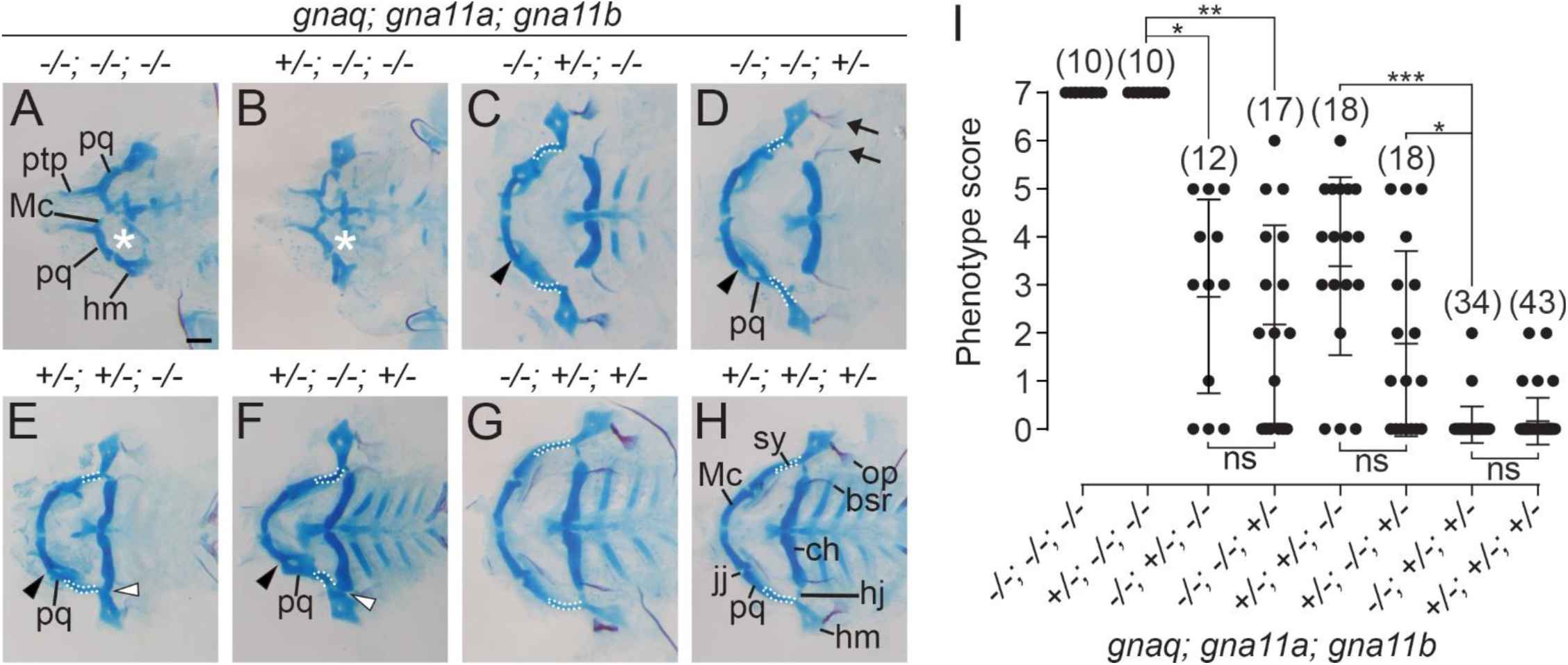
gna11a and *gna11b,* but not *gnaq,* are necessary for development of Edn1/Ednra-dependent structures. **(A-H)** Representative flat-mounts of the viscerocranium at 6 dpf, shown in ventral view, for selected allelic combinations. In C-H, white outlines highlight the symplectic. In A,B, the white asterisks indicate absence of the ceratohyal. In C-F, black arrowheads indicate fusion of the jaw joint. In E,F, the white arrowheads indicate fusion of the hyomandibular joint. **(I)** Overall severity of defects for genotypes shown in A-H were quantified as a “phenotype score”. Individual larvae were scored for the total number of Edn1/Ednra-dependent skeletal elements exhibiting defects (ignoring sidedness). A phenotype score of 0 correspond to a wild-type phenotype, and a score of 7 indicates all Edn1/Ednra-dependent structures on at least one side were affected. One dot represents an individual larva. The middle line is the mean. Error bar is standard error of the mean. The n values for respective genotypes are indicated in parentheses. Statistical significance was determined with Kruskal-Wallis test with Dunn’s multiple comparisons (*; *p* < 0.05, **; *p* < 0.01, ***; *p* < 0.001, ns; not significant). Scale bar is 100 μm.

### Gq is sufficient for lower jaw specification

To determine whether Gq/11 signaling activity is sufficient for lower jaw development, we tested if we could rescue the *edn1^-/-^* mutant phenotype by expressing a constitutively active form of Gq, Gq-Q209L (Fig. 6) (Kalinec et al., 1992). The Q209L amino acid substitution prevents GTP hydrolysis and maintains Gq in an active signaling state independent of receptor activity (Kleuss et al., 1994). Given the functional redundancy of Gq and G11 proteins (Malbon, 2005; Wettschureck and Offermanns, 2005), we surmised that Gq-Q209L would activate the same signaling pathways as zebrafish G11a and G11b proteins. In order to control the timing of induction for Gq activity, we made a transgenic line that expresses Gq-Q209L under the regulation of a *hsp70l* heat shock-inducible promoter (Halloran et al., 2000), *hsp70l:Gq-Q209L.* Embryos generated from *edn1^+/-^* and *edn1^+/-^;hsp70l:Gq-Q209L* mating pairs were heat-shocked at 16 hpf and then processed for fluorescence in situ hybridization at 28 hpf or skeletal preparations at 4 dpf. We performed skeletal preparations at 4 dpf instead of 6 dpf because a small percentage of *hsp70l:Gq-Q209L-*positive larvae exhibited early embryonic lethality starting at 5 dpf due to substantial cardiac edema (Fig. S9). In heat-shocked embryos, the majority of Edn1/Ednra-dependent structures were restored in 100% (9/9) of *edn1^-/-^;hsp70l:Gq-Q209L* larvae (Fig. 6D). The palatoquadrate, symplectic, ceratohyal, Meckel’s cartilage and the hyomandibular joint were restored at the highest frequency, whereas jaw joint restoration occurred at a lower frequency (Fig. 6E). We also observed apparent homeotic transformations of upper jaw structures to lower jaw-like structures in all transgene-positive larvae that were heat-shocked (Fig. 6F-H). In most larvae, the pterygoid process of the palatoquadrate, typically a short, pointed structure, was transformed into a long, broad piece of cartilage resembling Meckel’s cartilage. The dorsal portion of the hyomandibular cartilage was also hypoplastic and misshapen. Similar changes to the pterygoid process and hyomandibular cartilages have been reported in larvae with ectopic Ednra signaling activity in the dorsal domain of the first and second arches (Kimmel et al., 2007; Alexander et al., 2011; Zuniga et al., 2011; Barske et al., 2018).

**Figure 6.**
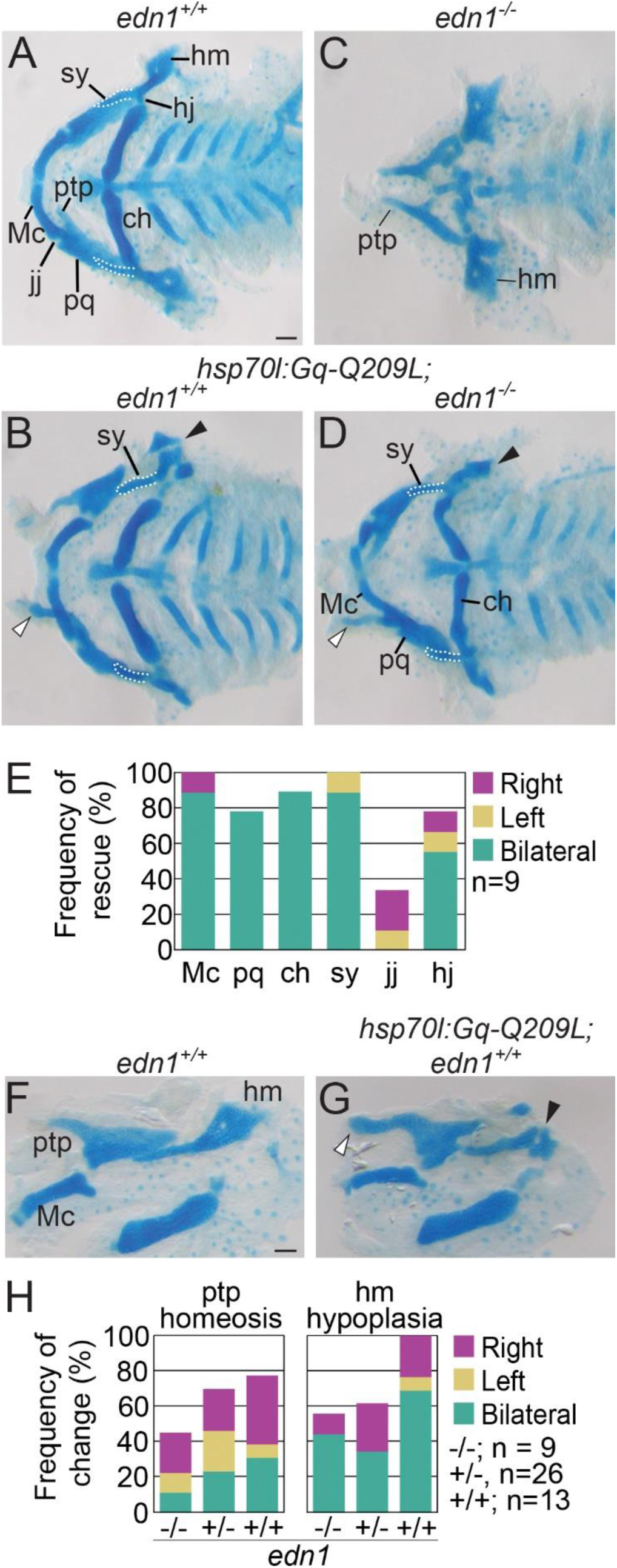
Induction of Gq activity rescues the *edn1^-/^*^-^ phenotype and causes ventralization of dorsal structures. **(A-D)** Flat-mounts of the viscerocranium at 4 dpf, shown in ventral view, of heat-shocked larvae. In *hsp70l:Gq-Q209L*;*edn1^+/+^*larvae (B), lower jaw structures were normal, though morphological changes were observed for the pterygoid process of the palatoquadrate (white arrowhead) and hyomandibular cartilage (black arrowhead). Compared to non-transgenic *edn1^-/-^*larvae (C), Edn1/Ednra-dependent skeletal elements were rescued in *hsp70l:Gq-Q209L*;*edn1^-/-^*larvae (D). Morphological changes in the pterygoid process of the palatoquadrate (white arrowhead) and hyomandibular cartilage (black arrowhead) were also observed in *hsp70l:Gq-Q209L edn1^-/-^* larvae (D) **(E)** Frequency of phenotype rescue for six Edn1/Ednra-dependent structures in heat-shocked *hsp70l:Gq-Q209L*:*edn1^-/-^* larvae (accounting for sidedness). Phenotype rescue was determined to be statistically significant with a chi-square test *(p=*0.0001), comparing the number of restored skeletal elements in non-transgenic *edn1^-/-^* larvae (0) to *hsp70l:Gq-Q209L*;*edn1^-/-^* larvae. **(F,G)** Flat-mounts of the viscerocranium at 4 dpf, in lateral view, of heat-shocked non-transgenic *edn1^+/+^* (F) and *hsp70l:Gq-Q209L; edn1^+/+^* larvae (G). Malformations of the pterygoid process of the palatoquadrate (white arrowhead) and hyomandibular cartilage (black arrowhead) are indicated in G. **(H)** Frequency of malformed pterygoid processes and hyomandibular cartilages (accounting for sidedness) in all *edn1* genotypes with the *hsp70l:Gq-Q209L* transgene. Scale bar is 100 μm.

These morphological changes were preceded by changes to patterning gene expression (Fig. 7). In 28 hpf *edn1^-/-^* embryos, the expression of *dlx5a,* a transcription factor essential for lower jaw specification, was severely reduced in the first and second pharyngeal arch mesenchyme (Fig. 7B). In *edn1^-/-^;hsp70l:Gq-Q209L* embryos that were heat-shocked, *dlx5a* expression was not only restored in the ventral and intermediate domains of the first and second arches, but also expanded into the dorsal domain (Fig. 7F). Similar expansion of *dlx5a* was observed in *edn1^+/+^;hsp70l:Gq-Q209L* embryos (Fig. 7E). Changes were also observed for *nr2f5*, an essential upper jaw specification factor (Barske et al., 2018) (Fig. 7C,D,G,H). In *edn1^+/+^* embryos, *nr2f5* expression was confined to the dorsal and intermediate domains of the first and second arches (Fig. 7C), while expression expanded into the ventral domains of the first and second arches in *edn1^-/-^* embryos (Fig. 7D). In *edn1^+/+^;hsp70l:Gq-Q209L* and *edn1^-/-^;hsp70l:Gq-Q209L* embryos, however, *nr2f5* was reduced overall (Fig. 7G,H). These data indicate that Gq is sufficient for induction of lower jaw patterning genes and subsequent development of lower jaw structures.

**Figure 7.**
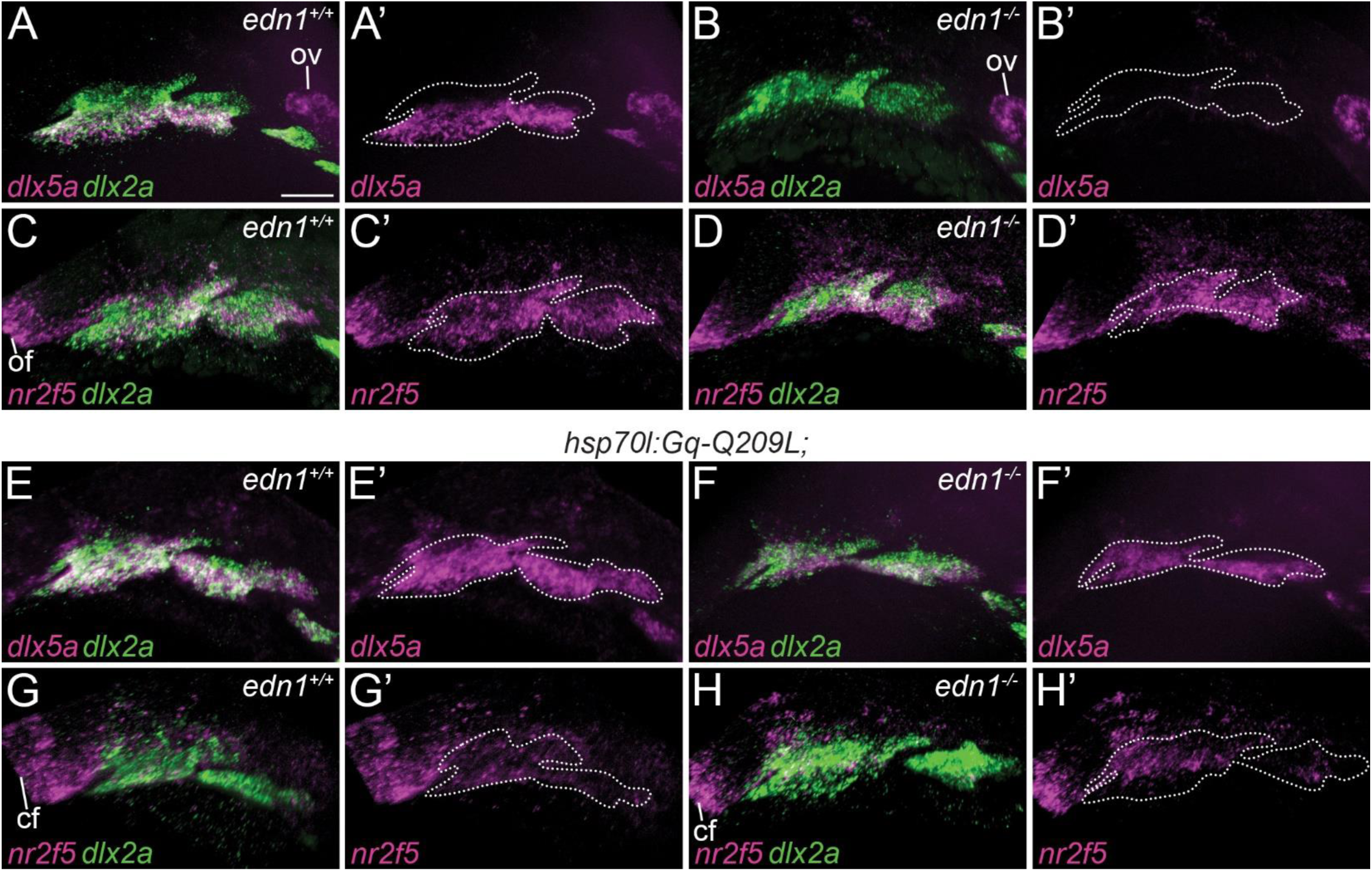
Induction of Gq activity upregulates *dlx5a* expression and downregulates *nr2f5* expression. The expression patterns of both *dlx5a* and *nr2f5*, both in magenta, are shown by two-color fluorescence in situ hybridization on 28 hpf embryos. Pharyngeal arches are labeled with *dlx2a,* in green. *dlx5a* and *dlx2a* (A,B,E,F) or *nr2f5* and *dlx2a* (C,D,G,H) are shown overlaid. *dlx5a* (A’,B’,E’,F’) and *nr2f5* (C’,D’,G’H’) are also shown alone, with the border of the pharyngeal arches indicated by white dashed lines. **(A,C)** Expression in non-transgenic *edn1^+/+^* embryos. **(E,G)** Expression in *hsp70l:Gq-Q209L; edn1^+/+^* embryos. **(B,D)** Expression in non-transgenic *edn1^-/-^* embryos. (**F,H)** Expression in *hsp70l:Gq-Q209L*:*edn1^-/-^* embryos. All embryos were heat-shocked. Images are representative of at least five embryos. Scale bar is 50 μm. cf; choroid fissure, ov; otic vesicle

## Discussion

In this study we addressed a long-standing question regarding the role of Gq/11 and potentially other Gα family members in lower jaw patterning (Sato et al., 2008a). Using the zebrafish model, we specifically interrogated the role of Gq/11 with three orthogonal approaches. We showed that inhibition of Gq/11 activity with a small molecule inhibitor or mutant alleles of *gnaq/gna11a/gna11b* produced craniofacial phenotypes similar to those observed in *edn1^-/-^*larvae and larvae injected with morpholinos targeting *ednraa and ednrab* (Miller et al., 2000; Nair et al., 2007), illustrating that Gq/11 is necessary for patterning of intermediate and ventral domains of the pharyngeal arches. We also showed that transgene-mediated expression of a constitutively active Gq protein, Gq-Q209L, can rescue lower jaw structures in *edn1^-/-^* embryos and can also transform the pterygoid process of the palatoquadrate to a Meckel’s cartilage-like structure, indicating that Gq/11 is sufficient to activate gene expression programs in cranial NCCs that establish lower jaw identity. Together, these results suggest that Gq/11 is the sole mediator of the Ednra signaling pathway in lower jaw patterning.

One of the most surprising findings from this work was the phenotypic discrepancy between zebrafish and mice lacking Gq and G11. The phenotype of *gnaq/gna11a/gna11b* mutant zebrafish larvae are identical to *edn1* mutants (Miller et al., 2000), with both exhibiting severe hypoplasia of the Meckel’s cartilage, fusion of the jaw joint and loss of the symplectic and ceratohyal cartilages. Similarly, the entire mandible of E18.5 *Edn1^-/-^*, *Ednra^-/-^* and *Ece1^-/-^* embryos undergoes a homeotic transformation into a maxilla-like structure (Kurihara et al., 1995; Clouthier et al., 1998; Yanagisawa et al., 1998b). In contrast, lower jaw defects in E18.5 *Gnaq^flox/flox^;Gna11^-/-^;P0-Cre* mouse embryos were limited to the proximal 2/3 of the mandible (Dettlaff-Swiercz et al., 2005) and more closely resembled zebrafish embryos treated with YM (Fig. 2,3) or reduced Gq/11 gene dosage (Fig. 5) than *Edn1^-/-^* or *Ednra^-/-^* embryos (Clouthier et al., 1998; Ozeki et al., 2004). One explanation for the partial lower jaw defect in *Gnaq^flox/flox^;Gna11^-/-^;P0-Cre* embryos is a technical issue related to P0-Cre. Gene recombination in first pharyngeal arch NCCs induced by P0-Cre appears less efficient compared to the NCC deletion strain *Wnt1-Cre* (Chen et al., 2017). Therefore, the residual levels of Gq expression and activity in cranial NCCs of *Gnaq^flox/flox^;Gna11^-/-^;P0-Cre* embryos may be sufficient to pattern the ventral domain but not the intermediate domain. This is consistent with numerous studies, including this one, that have reported that cranial NCCs in the intermediate domain are highly sensitive to perturbations that diminish Ednra signaling activity, such as treatment with YM (Fig. 2,3) or an Ednra antagonist (Ruest and Clouthier, 2009), reduction of Gq/11 gene dosage (Fig. 5) (Offermanns et al., 1998), or reduction of Edn1 levels with an Edn1 morpholino (Miller and Kimmel, 2001) or a mutant allele of *furina* (Walker et al., 2006), an enzyme in the Edn1 biosynthetic pathway. This sensitivity of cranial NCCs in the intermediate domain has been attributed to a morphogen gradient model for Edn1, which posits that extracellular concentrations of Edn1 are highest near the source, the ventral ectoderm, and diminishes towards the dorsal regions of the arch (Kimmel et al., 2003). This question regarding the role of Gq/11 in mouse lower jaw development could be resolved with *Wnt1-Cre*, which has been shown to induce sufficient levels of gene recombination in cranial NCCs in *Ednra^flox/flox^* embryos (*Ednra^flox/flox^;Wnt1-Cre*) to recapitulate the *Ednra^-/-^* phenotype (Ruest and Clouthier, 2009).

The zebrafish mutant alleles for Gq/11 genes revealed two additional differences between zebrafish and mice. First, mice and zebrafish embryos have different genetic requirements for Gq and G11 genes. In mice with conventional Gq/11 knockout alleles, embryos with one wild-type allele for *Gna11 (Gnaq^-/-^;Gna11^+/-^)* exhibited lower jaw defects, while embryos with one wild-type allele for *Gnaq* (*Gnaq^+/-^;Gna11^-/-^)* did not, indicating that *Gnaq* is primarily responsible for driving lower jaw patterning (Offermanns et al., 1998). In zebrafish, *gna11a* and *gna11b* were required for lower jaw patterning, while *gnaq* was dispensable (Fig. 5). The functional redundancy of Gq and G11 proteins (Figs. 5-7) (Malbon, 2005; Wettschureck and Offermanns, 2005) predicts that the signaling pathways activated downstream of Ednra in cranial NCCs of mice and zebrafish are likely the same, so what purpose is served by the differential usage of Gq/11 genes in zebrafish and mice? This difference may reflect divergent regulation of Gq/11 genes between teleosts and mammals. The teleost genome, which has undergone an additional round of duplication compared with mammals, exhibits significant divergence in the cis-regulatory elements (CRE) of duplicated genes and paralogs that result in different spatial and temporal expression patterns compared with other vertebrate species (Kassahn et al., 2009). Thus, the CREs for Gq/11 genes in teleosts may have also undergone changes that result in expression patterns that differ from mammals. This is consistent with our scRNA-seq analysis that shows *gnaq* expression is absent in cranial NCCs (Fig. S5), though the expression profile for *gnaq* must be experimentally verified. Thus, a comparative analysis of the CREs and expression profiles for Gq/11 genes in zebrafish and mice may provide insights into evolutionary mechanisms that drive changes to gene regulatory networks for cell signaling proteins, which could have implications for understanding how anatomical structures and functions have diverged between teleosts and mammals (Pires-daSilva and Sommer, 2003; de Mendoza et al., 2014).

The second difference was that zebrafish and mice are differentially sensitive to loss of Gq/11 signaling. *Gnaq^-/-^;Gna11^-/-^*mouse embryos exhibited mid-gestational lethality due in part to myocardial hypoplasia (Offermanns et al., 1998). However, zebrafish embryos that are triple homozygous for Gq/11 knockout alleles did not exhibit evidence of embryonic lethality prior to 6 dpf based on general morphology and normal Mendelian inheritance of alleles (Fig. S7). Although heart development was not examined in this study, it is possible that our triple homozygous Gq/11 mutant zebrafish have similar heart defects as mice given the conservation of Gq/11 function in craniofacial development. Unlike mouse embryos, zebrafish larvae can tolerate defective heart function up to 7 dpf due to a lack of dependence on oxygen from the circulatory system (Stainier et al., 1996). Therefore, this property of zebrafish physiology could be leveraged to examine the role of Gq/11 in heart development, which is not fully understood (Malbon, 2005; Wettschureck and Offermanns, 2005).

Our study also provides additional evidence for the utility and specificity of YM as a pharmacological tool to interrogate the role of Gq/11 in vivo (Shibata et al., 2022). Embryos treated with 100 uM YM produced phenotype and gene expression changes similar to embryos with partial loss of Edn1/Ednra signaling (Figs. 2, 3) (Miller and Kimmel, 2001; Walker et al., 2006; Nair et al., 2007). The window of sensitivity to YM also agrees with the window of sufficiency for endothelin signaling in lower jaw patterning from previous studies (Miller et al., 2000; Miller et al., 2003; Walker et al., 2006; Kimmel et al., 2007; Ruest and Clouthier, 2009; Vieux-Rochas et al., 2010; Alexander et al., 2011; Zuniga et al., 2011; Barske et al., 2016; Barske et al., 2018; Meinecke et al., 2018). Further, applying YM on *edn1* heterozygous mutants produced craniofacial phenotypes that were more severe in the spectrum of Edn1/Ednra-class phenotypes (Fig. 4). This additive effect of YM on sensitized backgrounds could be advantageous for studying other Gq/11-coupled receptors that might otherwise be embryonic lethal when knocked out. However, it is important to consider the system being studied, as we showed that the solubility limit of YM prevented compete inhibition of Gq/11 activity and thus full recapitulation of the *edn1* mutant phenotype. The effect of YM on zebrafish embryos is in stark contrast to the potency and efficacy of YM and its sister compound, FR900359, in cell culture assays, which exhibit an IC50 in the nanomolar concentration range (Nishimura et al., 2010; Schrage et al., 2015). The diminished inhibitory effect of YM in zebrafish embryos may be caused by poor perfusion into relevant tissues, inefficient cell permeability, or degradation. FR900359 exhibits slightly different physicochemical properties (Schlegel et al., 2021) and thus may exhibit greater effects on zebrafish embryos than YM (though FR900359 is not commercially available).

By establishing Gq/11 as the sole signaling mediator for Ednra in lower jaw patterning, we can begin to elucidate mechanisms of jaw patterning that remain poorly characterized. As demonstrated by our study and others, correct positioning of the upper and lower jaw patterning domains requires a specific balance of signaling inputs by Edn1/Ednra, Jagged/Notch and Nr2f nuclear receptors (Sato et al., 2008b; Zuniga et al., 2010; Alexander et al., 2011; Tavares and Clouthier, 2015; Barske et al., 2016; Barske et al., 2018; Sucharov et al., 2019; Kurihara et al., 2023). It remains unknown how these signaling pathways are integrated to elicit specific transcriptional outputs that refine the positional identities of cranial NCCs residing in and around the upper jaw-lower jaw boundary. Future studies will examine mechanisms of signal integration.

## Materials and methods

### Protein sequence alignment

Protein sequences were aligned using Clustal Omega (Madeira et al., 2019). The following are accession numbers for sequences used. For human proteins: Gq (NP_002063.2), G11 (NP_002058.2) and Gi1 (NP_002060.4). For zebrafish proteins: Gq (NP_001138271.1), G11a (NP_001038501.1), G11b (NP_001007774.1), G14 (NP_001003753.1), G14a (XP_683989.2), G15.1 (NP_001003626.2), G15.2 (XP_002667410.2), G15.3 (translated from mRNA sequence for *gna15.3*, XR_659583.3), G15.4 (NP_001038454.1), Gi1 (NP_957265.1), Gs (XP_001335732.1) and G12a (NP_001013295.1).

### Zebrafish strains and husbandry

All work was approved by the University of Colorado Institutional Animal Care and Use Committee (Protocol No. 00188). Zebrafish (*Danio rerio*) adults and embryos were raised and staged according to (Kimmel et al., 1995; Westerfield, 2007). Mutant and transgenic lines were maintained in the AB strain. The following mutant and transgenic lines were previously described: *sucker/edn1^tf216^* (Miller et al., 2000), *Tg(fli1a:EGFP)^γ1^* (Lawson and Weinstein, 2002) and *Tg(sox10:mRFP)^vu234^* (Kirby et al., 2006). In addition, five new lines were created: mutant alleles for *edn1^co3009^*, *gnaq^co3010^*, *gna11a^co3015^* and *gna11b^co3014^*, and the transgenic line *Tg*(*hsp70l:Gq-Q209L-IRES-EGFP,cmlc2:EGFP).* Construction of strains is described below.

#### Generation of deletion alleles

All mutant alleles were generated with the CRISPR/Cas9 gene-editing system as previously described (Hwang et al., 2013; Jao et al., 2013). Gene-specific target sequences were generated with CHOPCHOP v2 (Labun et al., 2016), and guide RNAs (gRNAs) were generated as previously described (Bassett et al., 2013). To generate *edn1^co3009^*, two target sequences were identified in exon 2. To generate *gnaq^co3010^* and *gna11b^co3014^,* target sequences were identified in exon 4 (Fig. S7). To generate *gna11a^co3015^,* a target sequence was identified in exon 3 (Fig. S7). The following are the target sequences used for each gene: *edn1;* 5’-GGAATAAGAGATGCTCCTGC-3’and 5’-GGACATAATATGGGTGAACA-3’*, gnaq;* 5’-GGCTGGGTGGGAATGTAGGAA-3’*, gna11a;* 5’-GGCGAGAGGTCGATGTCGAGA-3’*, gna11b;* 5’GGTGGGAAGGTACGAAGATT-3’. The individual target sequences were then incorporated into an oligo consisting of 5’-[T7 promoter sequence]-[Target sequence]-[Start of scaffold sequence]-3’. The T7 promoter sequence is 5’-AATTAATACGACTCACTATA-3’, and the start of the scaffold sequence is 5’-GTTTTAGAGCTAGAAATAGC-3’. DNA templates for gene-specific gRNAs were then generated by performing PCR with the target sequence-containing oligo and a separate oligo containing the gRNA scaffold sequence: 5’-GATCCGCACCGACTCGGTGCCACTTTTTCAAGTTGATAACGGACTAGCCTTATTTTAACTTGCTATTTCTAGCTCTAAAAC-3’. All oligos were purchased from IDT DNA Technologies. PCR reactions were performed with KAPA HiFi HotStart ReadyMix (Roche, 07958927001) (1x KAPA Taq, 500 nM target sequence oligo, 500 nM gRNA scaffold oligo) using the cycling parameters: 98°C for 30 seconds; 40 cycles of 98°C for 10 seconds, 60°C for 10 seconds, 72°C for 15 seconds; 72°C for 10 minutes. Following purification of the PCR product using the QIAquick PCR Purification Kit (Qiagen, 28104), the gRNA was synthesized using the MEGAscript T7 Transcription kit (Invitrogen, AM1334). Cas9 mRNA (codon-optimized for zebrafish) was generated as previously described (Jao et al., 2013). Briefly, pT3TS-nCas9n (Addgene plasmid # 46757) was linearized with *Xba*I restriction enzyme and then used as the template for in vitro transcription using the mMESSAGE mMACHINE T3 Transcription Kit (Invitrogen, AM1348).

Embryos were injected at the one-cell stage with 2 nl of solution containing 100 ng/ul Cas9 mRNA, 50 ng/ul gRNA and 0.025% phenol red (Sigma, P0290). To determine whether the gRNAs were excising the genomic loci of interest, at least 10 embryos per injected clutch were lysed in genomic DNA extraction buffer (10 mM Tris pH 8, 2 mM EDTA, 0.2% Triton X-100, 0.2 mg/ml Proteinase K) at 24 hpf, and the genomic loci of interest were PCR-amplified with genotyping primers. PCR amplicons were then analyzed by gel electrophoresis, with the presence of smears or ladders serving as indicators of potential genomic lesions. Clutches indicating genomic lesions were grown to adults (F0) and then crossed with wild-type ABs to generate F1 embryos. F1 adults were screened for genomic lesions at loci of interest by PCR, and the nature of the genomic lesions were characterized by Sanger sequencing. F1 adults harboring frameshift mutations were crossed with wild-type ABs to identify those that transmit mutant alleles to offspring in the expected Mendelian frequency. Germ line-stable F1 adults were kept and outcrossed to wild-type ABs to generate F2 fish used in this study. The following are genotyping primers used for each allele: *edn1^co3009^*; 5’- TAGGTGCTCCAGCATCTTTG-3’ and 5’-GGAGCGTTTCCAAGTCCATA-3’, *gnaq^co3010^;* 5’- TATGATGCCTTTTTGTCCACAG-3’ and 5’- CATTGTCAGACTCGACGAGAA-3’, *gna11a^co3015^*; 5’-TCAAGTCGTAAAATGGGTTGTG-3’ and 5’- TAAAAGCAGCAAATGACGACAC-3’, *gna11b^co3014^*; 5’- GAAGCATCCTTTACCAAACCAC-3’ and 5’- CGAACGAAGGCAGATGAATAAT-3’. The frameshift mutation in the *gna11b^co3014^* allele introduces a *Bce*AI restriction site. Thus, PCR reactions from the *gna11b^co3014^* genotyping assay were digested with *Bce*AI to further distinguish PCR amplicons from wild-type and mutant alleles.

#### Generation of transgenic line

The *hsp70l:Gq-Q209L-IRES-EGFP,cmlc2:EGFP* plasmid was generated using the Tol2kit as described in (Kwan et al., 2007). First, the middle entry vector containing Gq-Q209L, pME-Gq-Q209L, was generated. The Gq-Q209L cDNA was PCR-amplified from pc3.1-Gq-Q209L (Kanai et al., 2022) and then cloned into the pME-MCS vector using T4 ligase (New England Biolabs, M0202). pME-Gq-Q209L was then combined with p5E-hsp70l, p3E-IRES-GFPpA and pDestTol2CG2 using LR Clonase II Plus Enzyme Mix (Invitrogen,12538-120). To generate transposase mRNA, the pCS2FA-transposase plasmid was linearized with *Not*I restriction enzyme and then used as the template for in vitro transcription using the mMESSAGE mMACHINE SP6 Transcription Kit (Invitrogen, AM1340).

Embryos at the one-cell stage were injected with a 2 nl solution containing 12.5 ng/ul transposase mRNA, 12.5 ng/ul *hsp70l:Gq-Q209L-IRES-EGFP,cmlc2:EGFP* and 0.025% phenol red. Injected embryos were grown to adults (F0) and then crossed to wild-type ABs. Offspring (F1) were screened for the presence of the transgene by examining GFP expression in the heart. Transgene-positive, germ line-stable F1 fish were then crossed into *edn1^+/-^* fish to generate *Tg(hsp70l:GqQ209L-IRES-EGFP; edn1^+/-^* fish. The plasmids pME-MCS, p5E-hsp70l, p3E-IRES-GFPpA, pDestTol2CG2 and pCS2FA-transposase were provided by Kristen Kwan (Kwan et al., 2007).

### YM-254890 incubation

14 hpf embryos generated by intercrossing wild-type ABs were dechorionated in petri dishes coated with 0.5% agarose using E2 media (Westerfield, 2007) containing 2 mg/ml Pronase (Roche, 10165921001) at 25°C for 10-15 min. Embryos were subsequently washed in fresh E2 media in agarose-coated petri dishes three times. Dechorionated embryos in agarose-coated plates were then incubated with dimethyl sulfoxide (DMSO) or YM-254890 [Adipogen (AG-CN2-0509-M001) or Cayman Chemicals (29735)] dissolved in DMSO, from 16-36 hpf. At 36 hpf, embryos were washed with fresh E2 media three times and then processed for skeletal preparations at 6 dpf. 36 hpf embryos processed for fluorescence in situ hybridization with immunofluorescence were fixed overnight in 4% paraformaldehyde in phosphate buffered saline (PBS).

### Skeletal preparations

Larvae were stained with Alizarin Red and Alcian Blue as previously described (Walker and Kimmel, 2007; Brooks and Nichols, 2017).

### Single Cell RNA-sequencing

*Sample preparation for scRNA-seq.* Cranial neural crest cells were isolated from zebrafish embryos using a *fli1a:EGFP* and *sox10:mRFP* double labeling technique as previously described (Barske et al., 2016; Askary et al., 2017; Mitchell et al., 2021) with some modifications. In brief, 140 double positive embryos were dechorionated at 14 hpf with 2 mg/ml Pronase. 70 embryos were then treated with 100 uM YM-254890 or DMSO from 16-36 hpf. At 36 hpf, 10 embryos from each condition were set aside for skeletal preparations at 6 dpf. The heads of the remaining embryos were severed, pooled and deyolked by gentle pipetting in Ca^2+^-free Ringer’s solution (116 mM NaCl, 2.9 mM KCl, 5 mM HEPES, pH 7.0). The heads were then dissociated into single cells with mechanical agitation and enzymatic digestion for 15 minutes at 4°C with cold-activated protease from *Bacillus licheniformis (Sigma Aldrich, P5380)* in dissociation buffer (10 mg/ml *Bacillus licheniformis protease, 125 U/ml DNAse, 2.5 mM EDTA, PBS*), followed by enzyme neutralization with stop solution (30% fetal bovine serum, 0.8 mM CaCl_2_, in PBS). Cells were then centrifuged (400 x *g*), resuspended in cell suspension buffer [1% fetal bovine serum, 0.8 mM CaCl_2,_ Leibovitz’s L-15 Medium (Gibco, 21083-027)], filtered through a 70 uM strainer (PluriSelect, 43-10040-40), centrifuged again (400 x *g*) and resuspended in sorting buffer (1% fetal bovine serum, 1 mM EDTA, 25 mM HEPES). EGFP and mRFP double positive cells were then purified with fluorescence activated cell sorting using a MoFlo XDP100 (Beckman Coulter). A total of 72,000 and 86,000 double positive cells were purified for DMSO and YM-treated samples, respectively. Cells were once again centrifuged (400 x *g*) and resuspended in cell suspension buffer. After confirming cell viability with trypan blue (Gibco, 15250-061), 8000 cells for each condition were submitted to the University of Colorado Anschutz Medical Campus Genomics and Microarray Core for single cell RNA-sequencing. Singe cell capture, barcoding and library generation was performed using 10X Genomics single cell RNA-sequencing technology (Chromium Controller and Next GEM single Cell 3’ Kit v3.1, Dual Index). Libraries were processed for paired-end sequencing with a read depth of 75,000/cell using NovaSeq6000 (Illumina).

### scRNAseq analysis

Sequence reads were aligned to the *Danio rerio* reference genome (GRCz11) and converted to count matrices with Cell Ranger v5.0.1 (10X Genomics). Subsequent analyses of count matrices were performed with Seurat version 4.3.0 in R studio (Hao et al., 2021). Individual Seurat objects were created for DMSO and YM-treated samples, and cells in each dataset were filtered for quality. For DMSO-treated samples, 3700 cells were obtained after filtering for cells with 200-3000 unique feature counts and less than 5% mitochondrial DNA. For YM-treated cells, 4506 cells were obtained after filtering for cells with 200-3500 unique feature counts and less than 5% mitochondrial DNA. Datasets for DMSO or YM-treated samples were then merged, normalized and scaled, with cell cycle phase-associated genes regressed out. A list of S- and G2M-phase genes built into Seurat was converted to zebrafish identifiers using BioMart and then used for cell cycle regression (Nestorowa et al., 2016). Converted gene lists were manually checked and edited for any changes in nomenclature/duplications between the reference and Ensembl database. Linear dimensional reduction was then performed with principal component analysis (PCA), and the top 30 PCAs were used to generate and visualize clusters with Uniform Manifold Approximation and Projection (UMAP) nonlinear dimensional reduction technique. Marker genes for clusters were obtained with a Wilcoxon Rank Sum test, and cluster identities were determined using published, experimentally verified marker genes (Nair et al., 2007; Talbot et al., 2010; Barske et al., 2016; Askary et al., 2017; Barske et al., 2018; Fabian et al., 2022). The approximate population of cells representing the intermediate domain were determined with a feature plot encompassing known intermediate domain marker genes, *ccn2b, dlx3b*, *dlx4a*, *dlx4b*, *emx2*, *fgfbp2a*, *foxc1b, foxd1, fsta, grem2b, igfbp5b, msx1a, nkx3.2* and *shox* (Fig. S4). Differential expression (DE) analysis was performed using Wilcoxon Rank Sum test between equivalent clusters of cells in DMSO and YM-treated samples.

### In situ hybridization and immunofluorescence

Two-color fluorescence in situ hybridization was performed as previously described (Talbot et al., 2010). Fluorescence in situ hybridization with immunofluorescence was performed as previously described (Mitchell et al., 2021). All probes have been previously described: *dlx2a* (Akimenko et al., 1994)*, dlx4b* (Ellies et al., 1997)*, dlx5a* (Walker et al., 2006) and *nr2f5* (Barske et al., 2018). The plasmid for *nr2f5* was provided by Lindsey Barske.

### Microscopy

Embryos processed for fluorescence in situ hybridization and immunofluorescence were mounted in 0.4% agarose in PBS on a glass-bottom dish (MatTek, P35G-1.5-10-C). All images were taken using a 20x air objective on a DMi8 microscope (Leica) equipped with an Andor Dragonfly confocal unit (Oxford Instruments). Images were processed using the Imaris image analysis software (Oxford Instruments). Skeletal preparations were imaged on a SZX12 stereo microscope (Olympus) equipped with a SC100 camera (Olympus).

### Heat shock protocol

Embryos generated from crossing *Tg(hsp70l:GqQ209L-IRES-GFP,cmlc2:EGFP);edn1^+/-^*and *edn1^+/-^* adults were heat-shocked at 16 hpf for 10 minutes in a 38°C water bath and then returned to 28.5°C. Transgenic and non-transgenic embryos were separated based on cardiac EGFP expression and monitored daily until 4 dpf to account for embryos developing cardiac edema, and then processed for skeletal preparations. Embryos processed for in situ hybridization were fixed overnight in 4% paraformaldehyde in PBS at 28 hpf.

### Statistical analysis

Statistical analyses for Kruskal-Wallis test (nonparametric one-way ANOVA) with Dunn’s multiple comparisons test and chi-square tests were performed in Prism (GraphPad).

## Acknowledgements

We thank Christine Archer, Ainsley Gilbard and Olivia Gomez for zebrafish care, the Genomics and Flow Cytometry cores at the University of Colorado Anschutz Medical Campus for assistance with sample preparation for scRNA-sequencing, members of the Nichols lab and the craniofacial biology interest group (“11 at 11” joint lab meetings) for helpful comments and feedback and Dr. Joan Hooper for sharing Seurat code.

## Author contributions

Conceptualization: S.M.K., D.E.C., J.T.N; Methodology: S.M.K., D.E.C., J.T.N.; Software: S.M.K., E.S.L.; Formal analysis: S.M.K., E.S.L.; Investigation: S.M.K., C.R.G., M.R.A., S.A.S., E.P.B., J.S.; Resources: S.M.K., D.E.C., J.T.N.; Writing – original draft: S.M.K., D.E.C.; Writing – review & editing: S.M.K., D.E.C., J.T.N., E.S.L.; Visualization: S.M.K., E.S.L.; Supervision: S.M.K., D.E.C., J.T.N.; Project administration: S.M.K., D.E.C., J.T.N.; Funding acquisition: S.M.K., D.E.C., J.T.N.

## Data availability

The raw and processed single cell RNA-sequencing datasets are available in the Gene Expression Omnibus (GEO) database under accession number: GSE275013.

## Supplementary Information

**Fig. S1.**
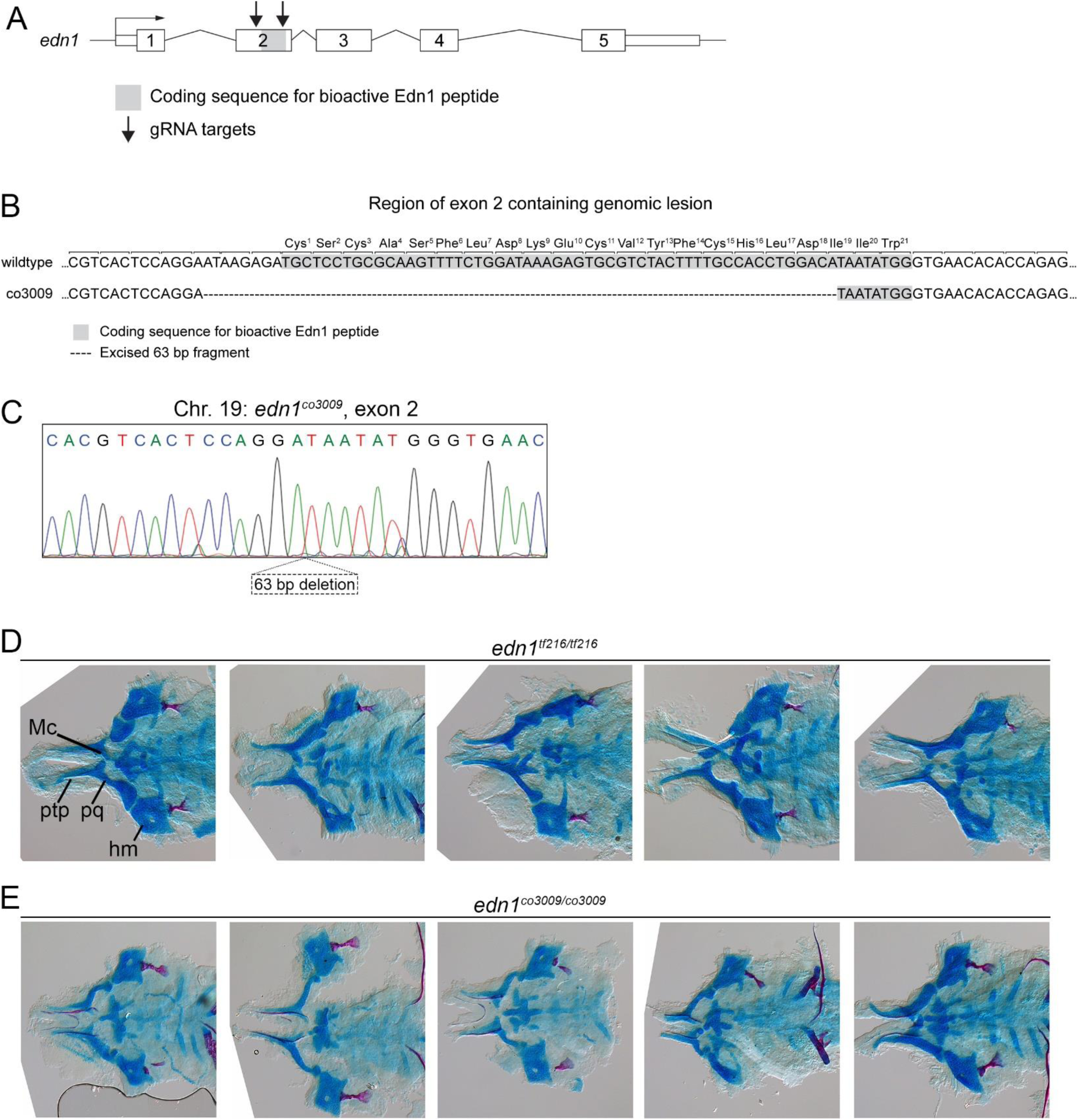
Targeting strategy for new *edn1* allele and characterization of genomic feature and phenotype. **(A)** Schematic of gene locus for *edn1* and targeting sites for sgRNAs (arrows). The coding sequence for the bioactive portion of Edn1 is in exon2 (in grey). **(B)** The co3009 allele contains a 63 bp in-frame deletion in exon 2, excising a majority of the coding sequence for the bioactive peptide. Excised nucleotides are indicated with dashed lines. The amino acids for the bioactive Edn1 peptide are labeled. The open reading frame is indicated with brackets. **(C)** Sanger sequencing trace confirms the genomic lesion. **(D,E)** Craniofacial phenotypes are indistinguishable between larvae homozygous for the *tf216/sucker* allele and the *co3009* deletion allele. Five representative flat-mounts of the viscerocranium at 6 dpf are shown for (D) *edn1^tf216/tf216^* (n=63) and (E) *edn1^co3009/co3009^* (n=41) larvae.

**Fig. S2.**
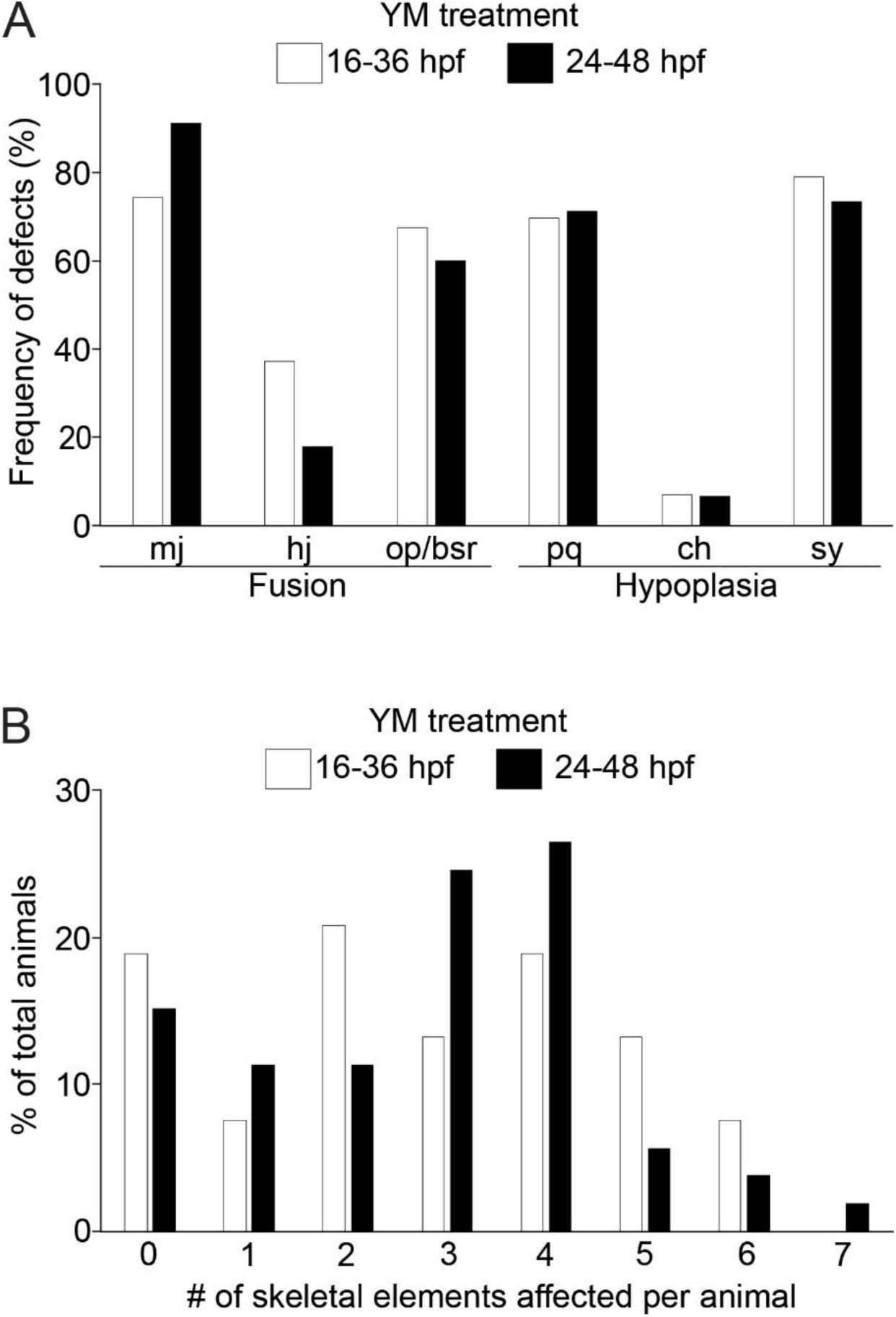
Frequency and overall severity of defects in YM-treated larvae. Shown are results for embryos treated with YM between 16-36 hpf vs 24-48 hpf. **(A)** Frequency of defects in specific skeletal elements for all larvae. (*p=*0.1, chi-square test) **(B)** Overall severity of defects, expressed as the number of skeletal elements affected per individual larva.

**Fig. S3.**
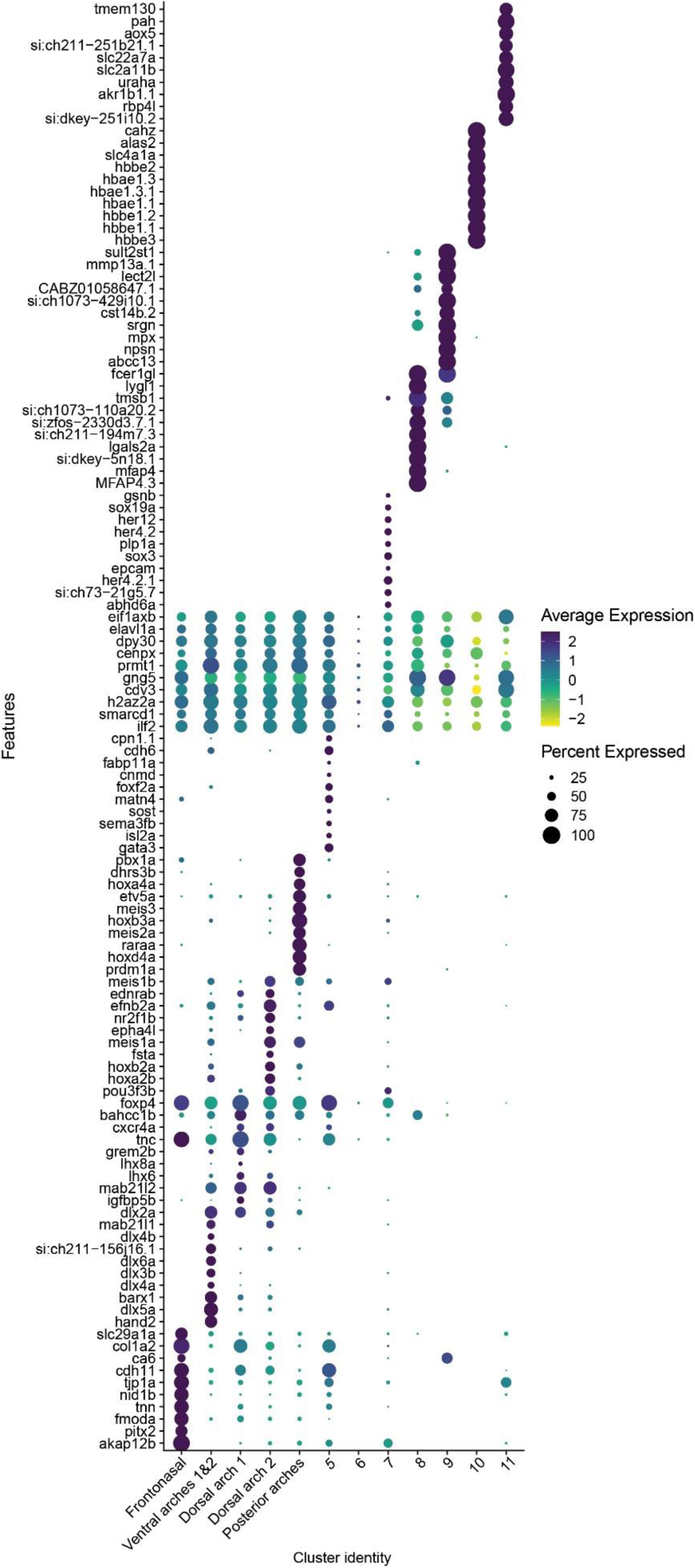
Dot plot analysis of marker genes. Shown are top ten genes per cluster based on adjusted p value.

**Fig. S4.**
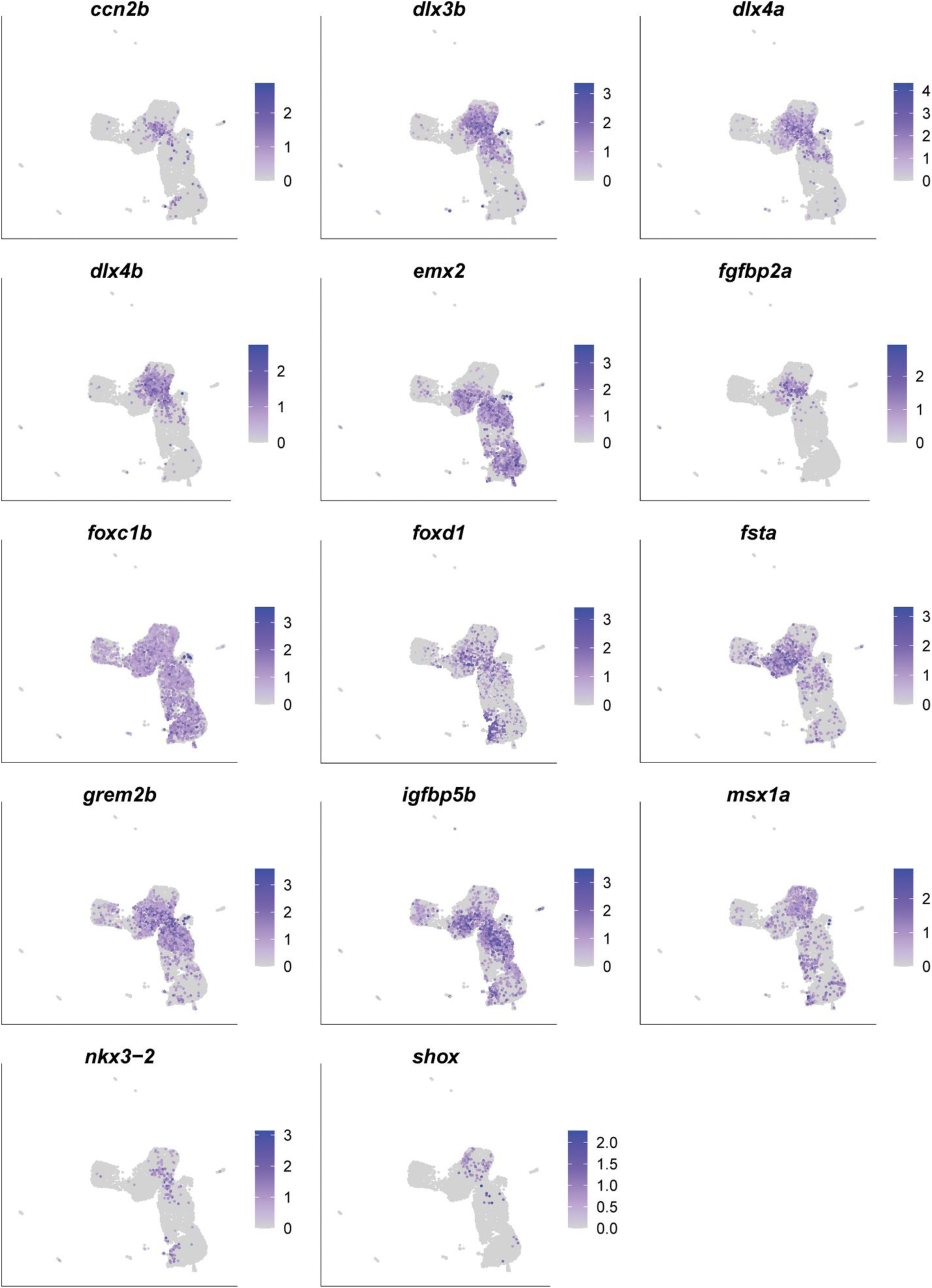
Feature maps of marker genes used to define the intermediate domain. The selected marker genes have been experimentally verified. Scale is average expression.

**Fig. S5.**
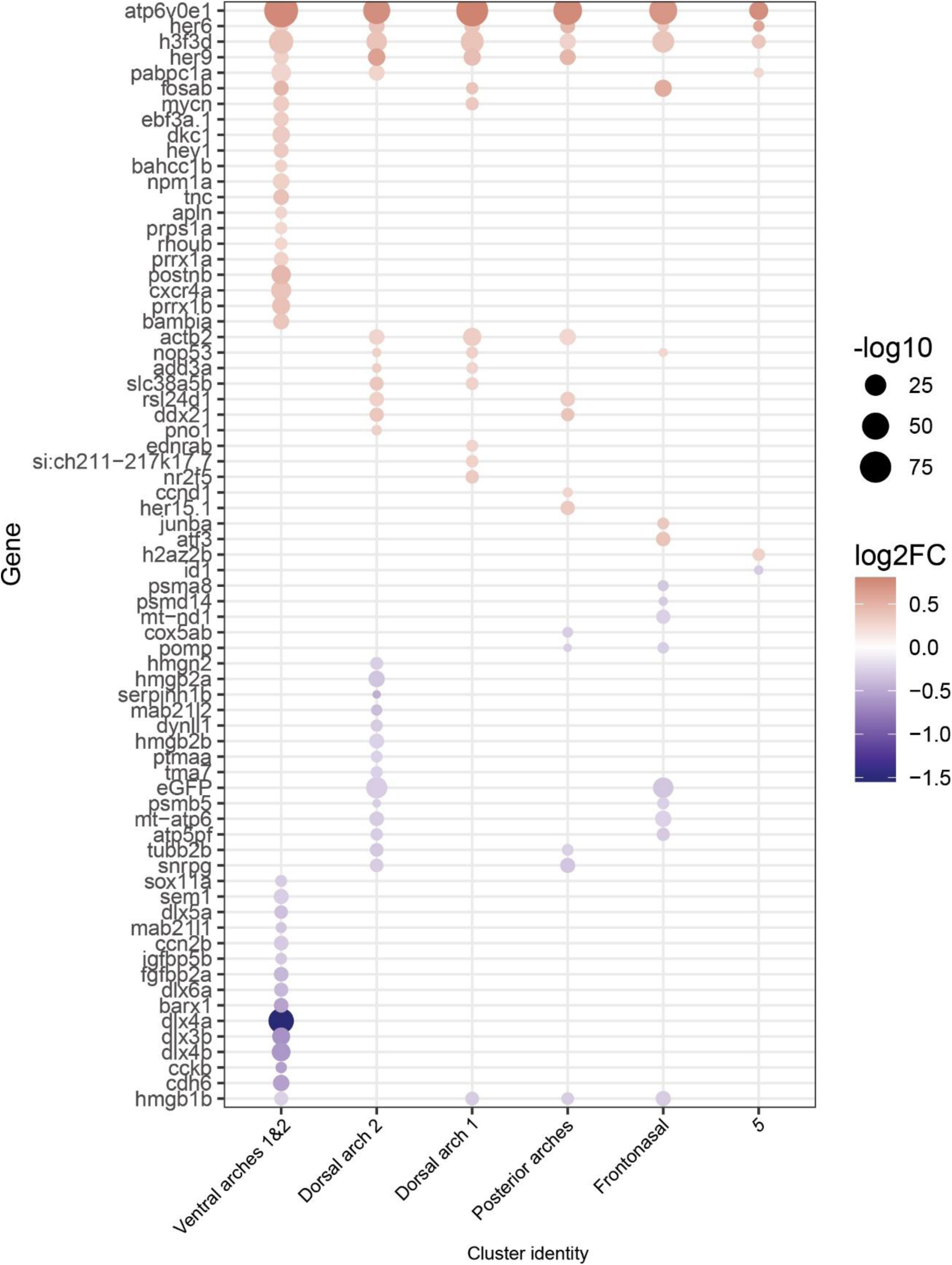
Dot plot of differentially expressed genes. All differentially expressed genes based on adjusted p values in YM-treated samples relative to DMSO-treated control samples, across equivalent clusters. Differential expression is displayed as log2-fold change (scale bar).

**Fig. S6.**
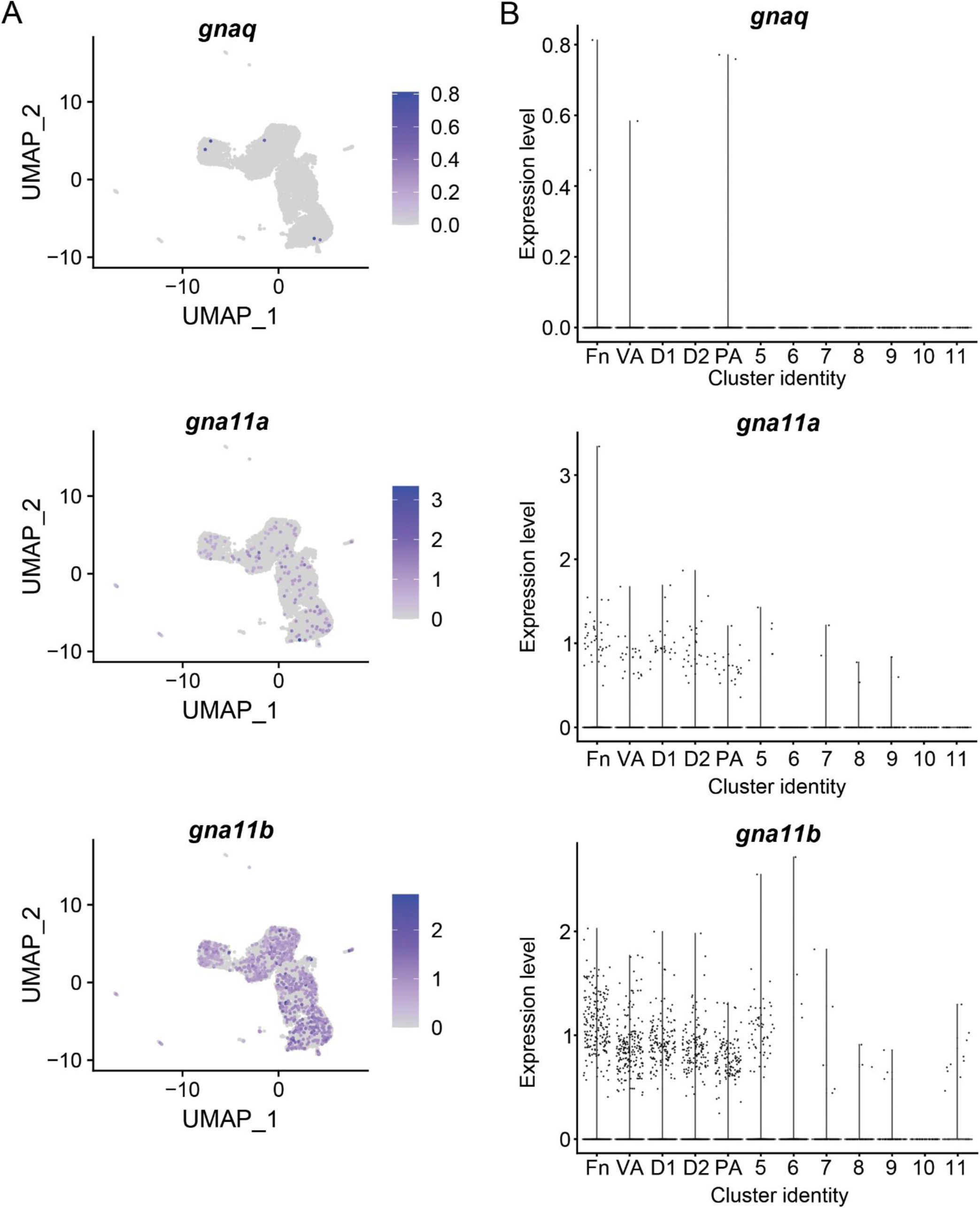
*gnaq, gna11a* and *gna11b* are differentially expressed in cranial neural crest cells. Expression profiles for *gnaq, gna11a* and *gna11b* are shown in **(A)** UMAP plots and **(B)** Violin plots. Scale is average expression. D1; Dorsal arch 1, D2; Dorsal arch 2, Fn; Frontonasal, PA; Posterior arches, VA; Ventral arches 1&2

**Fig. S7.**
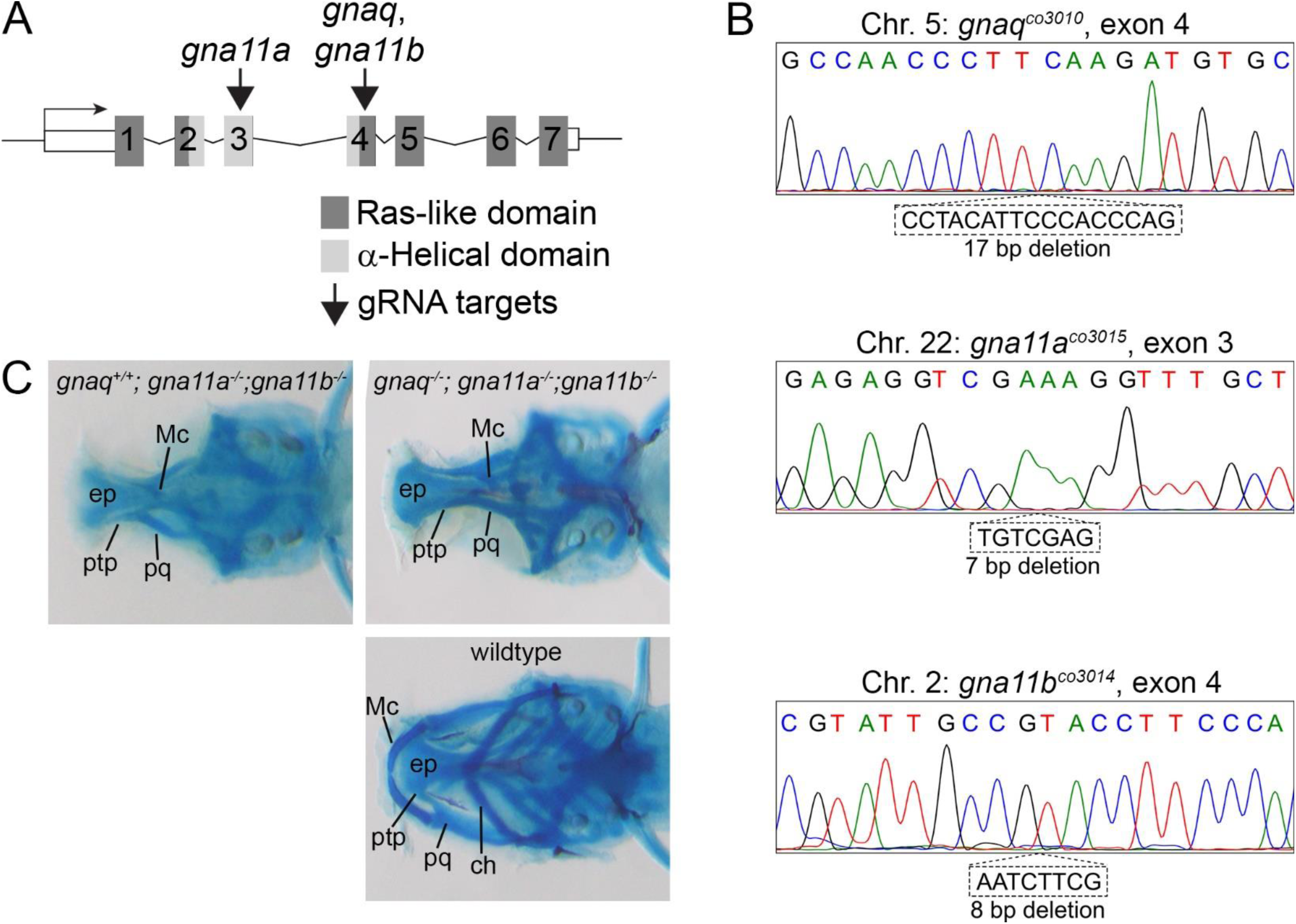
Targeting strategy for CRISPR/Cas9 gene-editing and characterized alleles. **(A)** Schematic of a generalized gene locus for *gnaq, gna11a* and *gna11b*, and the targeting sites for sgRNAs. Numbers indicate exons. Exons encoding for the Ras-like and α-Helical domains are indicated. **(B)** Sanger sequencing traces confirm frameshift mutations for the alleles *gnaq^co3010^*, *gna11a^co3015^* and *gna11b^co3014^.* For all alleles, the respective deletions result in a premature stop codon in exon 4. **(C)** Representative phenotypes for *gnaq^+/+^;gna11a^-/-^;gna11b^-/-^*, *gnaq^-/-^;gna11a^-/-^;gna11b^-/-^* and wildtype. Shown are ventral views of whole-mount skeletal preparations for 6 dpf larvae, with eyes removed. The ethmoid plate (ep) is a structure of the neurocranium. ch; ceratohyal, Mc; Meckel’s cartilage, pq; palatoquadrate, ptp; pterygoid process of the palatoquadrate

**Fig. S8.**
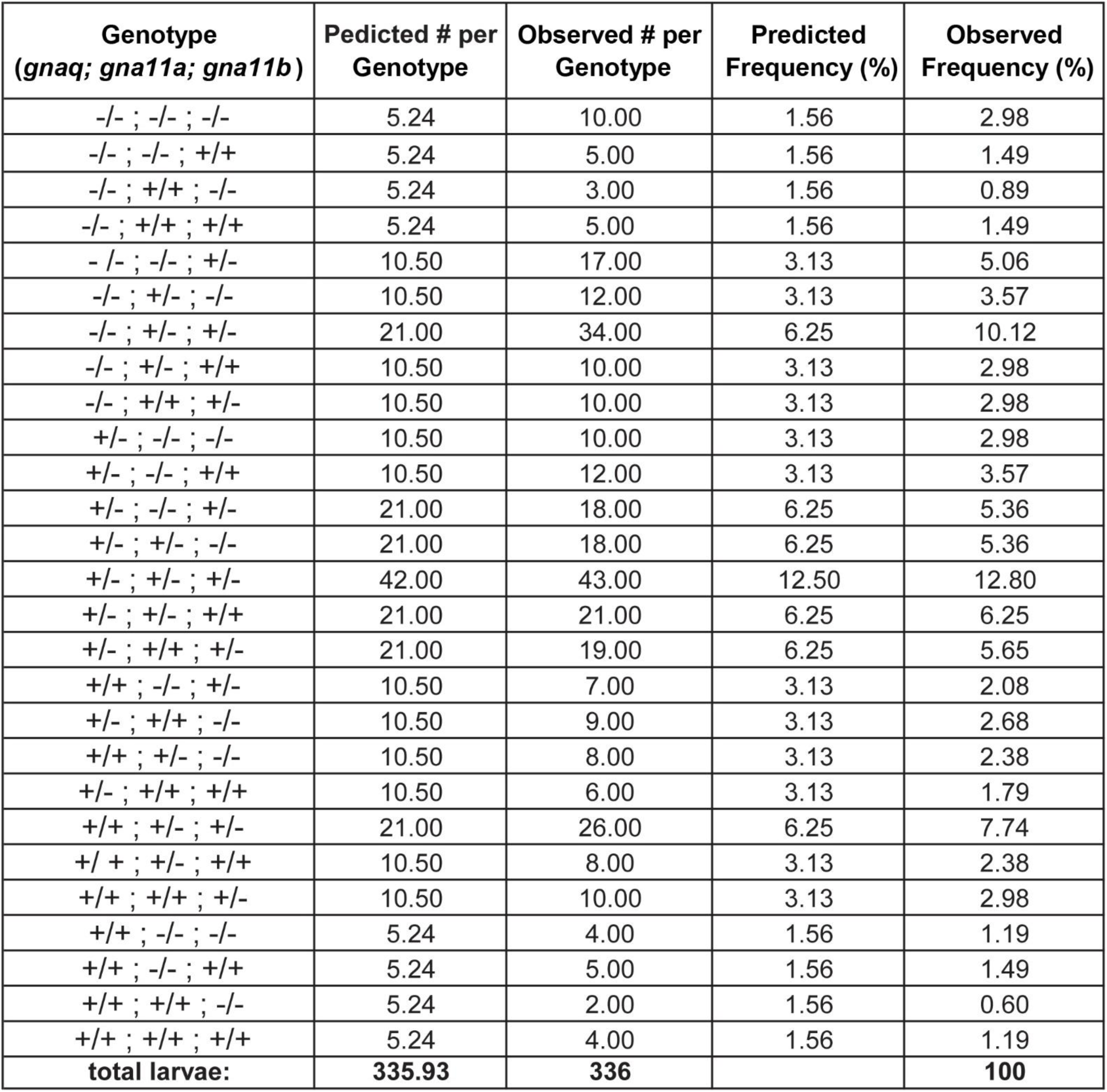
Table of predicted and observed genotypes obtained from crossing *gnaq^+/-^;gna11a^+/-^;gna11b^+/-^* animals. Values represent the sum of three independent clutches. Differences in predicted and observed values are not statistically significant (*p*=0.4, chi-square test)

**Fig. S9.**
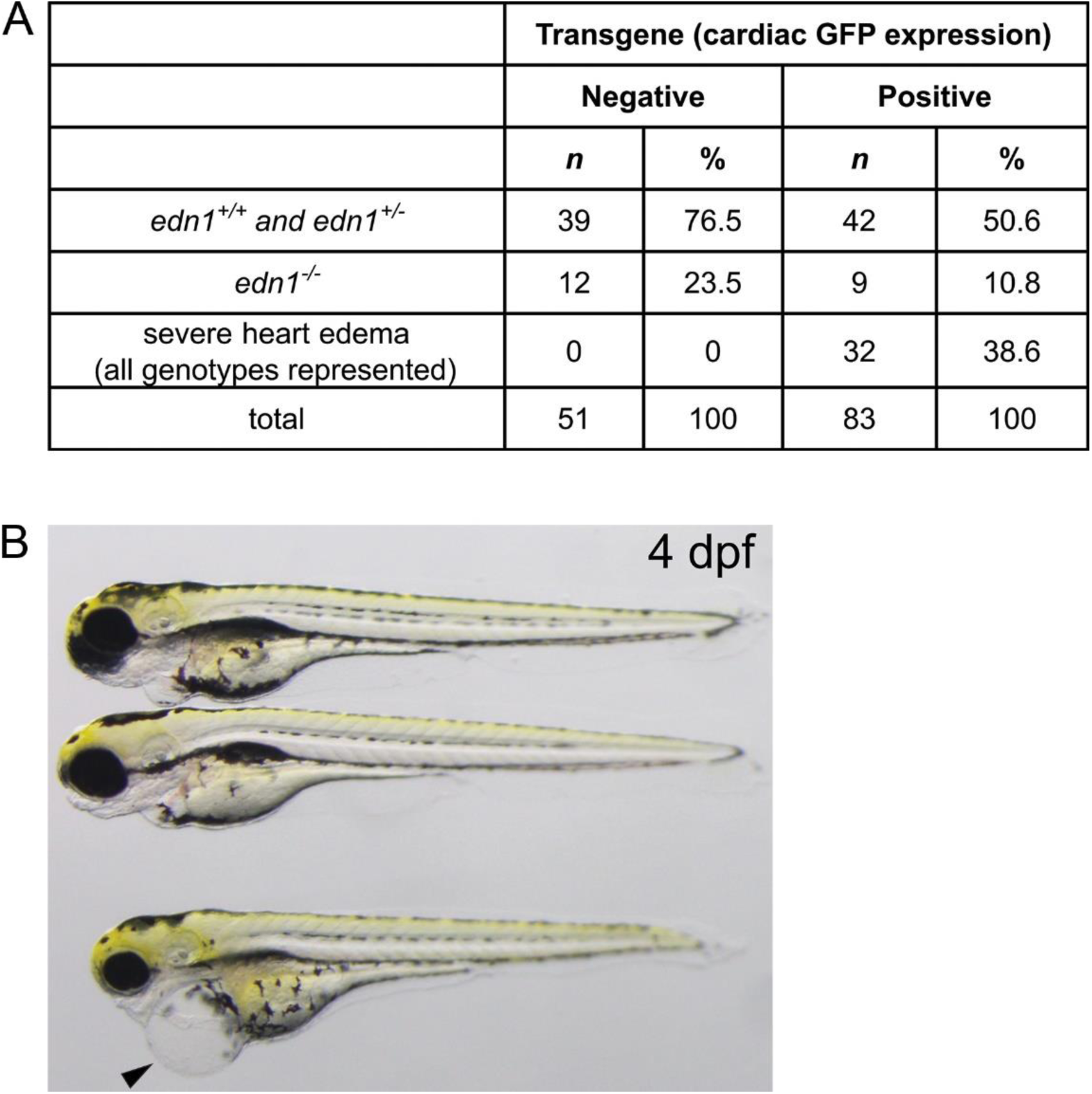
Frequency and example of heart edema associated with the *hsp70l:Gq-Q209L* transgene. **(A)** Table showing the frequency of genotypes and heart edema observed from embryos generated from *edn1^+/-^* x *edn1^+/-^;hsp70l:Gq-Q209L* crosses, which were subsequently heat-shocked. Heart edema was associated only with transgene-positive larvae, for all *edn1* genotypes (*edn1^+/+^, edn1^+/-^, edn1^-/-^)*. Larvae exhibiting heart edema (n=32) were excluded from the analysis. **(B)** Example of severe heart edema (arrow) associated with the *hsp70l:Gq-Q209L* transgene.

